# Pan-cancer detection of driver genes at the single-patient resolution

**DOI:** 10.1101/2020.06.12.147983

**Authors:** Joel Nulsen, Hrvoje Misetic, Christopher Yau, Francesca D. Ciccarelli

## Abstract

**Background:** Identifying the complete repertoire of genes that drive cancer in individual patients is crucial for precision oncology. Most established methods identify driver genes that are recurrently altered across patient cohorts. However, mapping these genes back to patients leaves a sizeable fraction with few or no drivers, hindering our understanding of cancer mechanisms and limiting the choice of therapeutic interventions.

**Results:** We present sysSVM2, a machine learning software that integrates cancer genetic alterations with gene systems-level properties to predict drivers in individual patients. Using simulated pan-cancer data, we optimise sysSVM2 for application to any cancer type. We benchmark its performance on real cancer data and validate its applicability to a rare cancer type with few known driver genes. We show that drivers predicted by sysSVM2 have a low false-positive rate, are stable and disrupt well-known cancer-related pathways.

**Conclusions:** sysSVM2 can be used to identify driver alterations in patients lacking sufficient canonical drivers or belonging to rare cancer types for which assembling a large enough cohort is challenging, furthering the goals of precision oncology. As resources for the community, we provide the code to implement sysSVM2 and the pre-trained models in all TCGA cancer types (https://github.com/ciccalab/sysSVM2).

## BACKGROUND

Cancer is characterised by the acquisition of somatic alterations of the genome, the majority of which are thought to have little or no phenotypic consequence for the development of the disease. Identifying the genes whose alterations instead have a role in driving cancer (cancer drivers) is one of the major goals of cancer genomics and numerous methods have been developed so far to achieve this.

Most of these methods work at the cohort-level, which means that they identify driver genes within a cohort of patients. For example, recurrence-based methods such as MutSigCV (1) and MuSiC (2) search for genes whose mutation rate (single nucleotide variants (SNVs) and small insertions or deletions (indels) per nucleotide) is above the background level. This is because mutations in cancer drivers are more likely to become fixed and recur across samples than those in non-driver genes. GISTIC2 (3) adopts a similar approach for recurrent copy number variants (CNVs). OncodriveCLUST (4) and ActiveDriver (5) look specifically for mutations clustering in hotspot positions or encoding post-translational modification sites. TUSON (6) and 20/20+ (7) predict new drivers based on features of canonical oncogenes and tumour suppressors, including the proportion of missense or loss-of-function to silent mutations occurring across patients. dNdScv (8) computes the nonsilent to silent mutation ratio to identify gene mutations under positive selection, while OncodriveFM (9) focuses on biases towards variants of high functional impact. Finally, network-based methods like HotNet2 (10) incorporate gene interaction networks to identify significantly altered modules of genes within the cohort. Albeit with different approaches, all these methods rely on the comparison of alterations and/or altered genes across patients.

Cohort-level methods have been of great value leading to the identification of more than 2,000 well-established (canonical) or candidate cancer driver genes (11, 12). However, these approaches fail to identify rare driver events that occur in small cohorts or even in single patients because of low statistical power. Moreover, they are not ideal for application in the clinical setting because they return lists of drivers in entire cohorts, rather than predictions in individual patients.

Patient-level methods ideally predict cancer drivers in each patient but are more challenging to implement. A few attempts such as OncoIMPACT (13), DriverNet (14) and DawnRank (15) combine transcriptomic and genomic data to identify gene network deregulations in individual samples. Such methods require user-specified gene networks and deregulation thresholds, which can affect their results (13). In addition, matched exome and transcriptome data from the same sample are not always available, especially in clinical settings where shotgun transcriptomic sequencing is still rare. Alternative approaches such as PHIAL (16) match the patient mutations with databases of known clinically actionable or driver alterations but have a limited capacity to identify as-yet unknown driver alterations. To overcome this limitation, iCAGES (17) combines deleteriousness predictions and curated database annotations to learn features of true positive and true negative driver alterations.

We recently developed sysSVM, a patient-level driver detection method based on one-class support vector machines (SVMs) (18). sysSVM learns the distinct molecular features (damaging somatic alterations) and systems-level features (gene properties) of canonical drivers. It then predicts as drivers the altered genes in individual patients that best resemble these features. When applied to 261 patients with oesophageal adenocarcinomas, sysSVM successfully identified the driver events in every patient (18).

Here, we further develop sysSVM to be applied to any cancer type and benchmark it against other available approaches, showing that it has a lower false positive rate and better patient coverage. We also develop optimal models for identifying driver genes in all 34 cancer types available in The Cancer Genome Atlas (TCGA) (19) and validate them in osteosarcoma, a rare cancer type that was not part of TCGA. The software, optimised models and their associated driver predictions are provided as a resource that can be used to identify and study driver events in cancers at the single patient resolution.

## IMPLEMENTATION

The sysSVM approach to driver detection prioritises genes with features similar to those of canonical cancer drivers, *i.e.* genes whose modifications have experimentally proven roles in cancer initiation and progression (Supplementary Note, Additional File 1). Canonical drivers differ from other human genes by an array of systems-level properties that define them as a group and do not strictly depend on the function of the single gene. These properties include gene duplicability in the human genome (20) and through vertebrate whole-genome duplications (21); gene essentiality across cell lines (11); breadth of expression in healthy tissues at the gene and protein levels (11, 22, 23); protein connectivity and global topology in the protein-protein interaction network (20); participation in protein complexes (22); number of targeting miRNAs (21); gene evolutionary origin (21); and protein length and domain organisation (22, 23) (Supplementary Table 1, Additional File 2). Canonical drivers can also be described using molecular properties that reflect the somatic alterations that they acquire in cancer. These include alterations with predicted damaging effects on protein function (copy number gains and losses as well as truncating, non-truncating damaging and hotspot mutations) and overall mutational burden and copy number of the gene (Supplementary Table 1, Additional File 2).

To leverage the systems-level and molecular properties of canonical drivers, sysSVM first identifies a set of true positive canonical drivers damaged within a cohort of patients (Figure 1A). It then uses the features of this positive set to train one-class SVMs based on four kernels (linear, radial, sigmoid, polynomial). Finally, it ranks the remaining damaged genes in individual cancer patients with a combined score that weights the kernels based on their sensitivity (Supplementary Note, Additional File 1). Highly ranked genes have the most similar properties to those of canonical drivers and will be then considered the cancer drivers for that patient. We use one-class SVMs for sysSVM because, while canonical drivers represent a reliable set of true positives, identifying a true negative set of non-cancer genes is not possible. For example, possible negative genes could be known false positives of driver gene detection methods (1, 22). However, these genes are representative of false positives rather than true negatives, so training a classifier on them is likely to introduce unwanted bias. A one-class support vector machine for novelty detection is therefore an optimal way to solve this issue.

**Figure 1.**
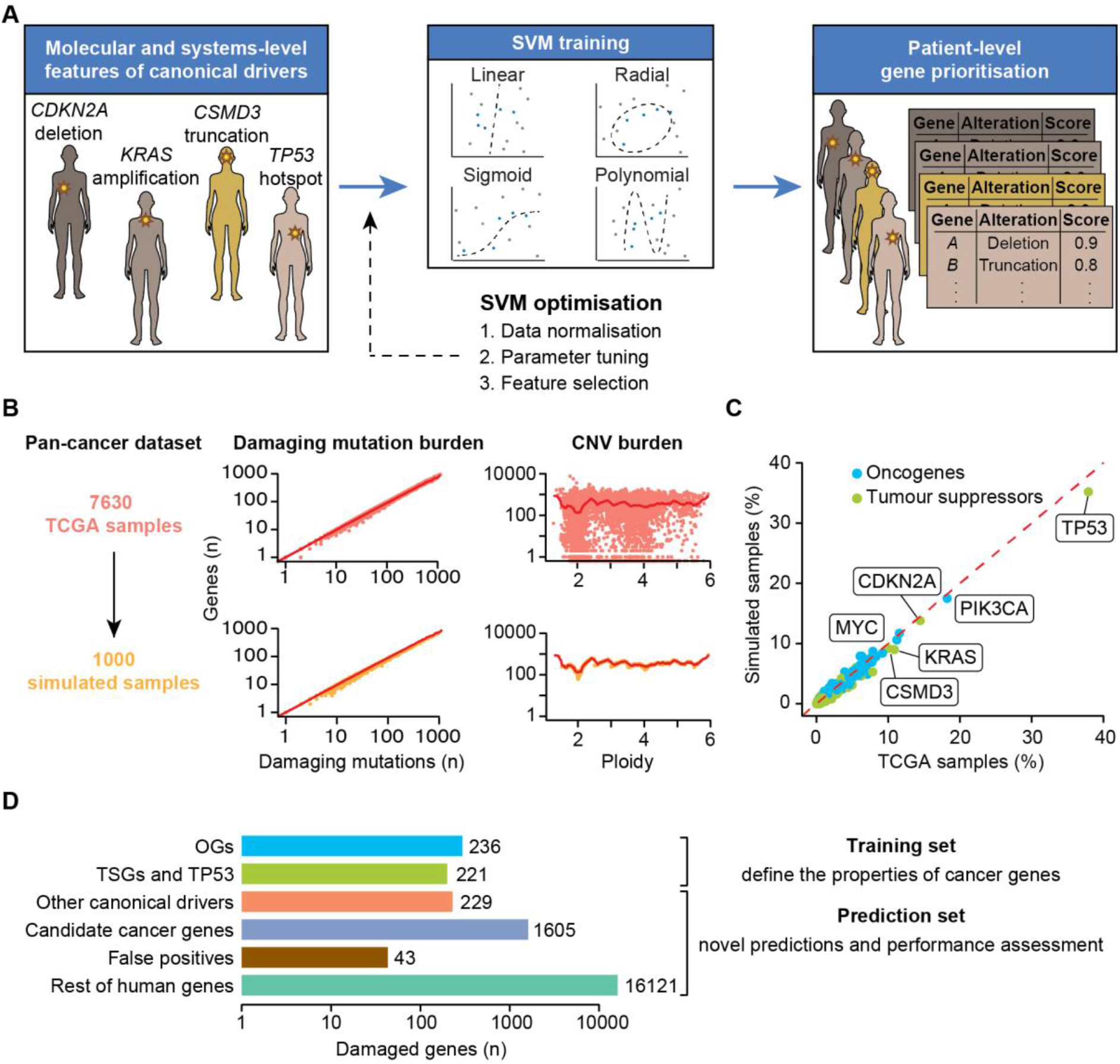
sysSVM approach for driver prioritisation. **A.** Overview of sysSVM. Molecular (somatic SNVs, indels and mutation burden) and systems-level features (Supplementary Table 1, Additional File 2) of damaged canonical drivers in the analysed samples are used for training. The best models of support vector machines (SVMs) with four kernels are selected using cross-validation and trained on the whole set of damaged canonical drivers. Finally, a combined score is used to prioritise driver genes in individual patients. The SVM implementation was generalised for optimal performance on a simulated cancer-agnostic dataset through data normalisation, parameter tuning and feature selection. **B.** Generation of a simulated reference cohort from TCGA data. Values of damaging mutation burden and ploidy were randomly assigned to samples. Damaged genes were then extracted from real samples with similar values of damaging mutation burden (+/−10% for mutations) and ploidy (+/−0.1 for CNVs). Dots represent individual TCGA (orange) or simulated (yellow) samples. Red lines indicate average numbers of genes with damaging mutations or CNVs in TCGA samples, for each given values of damaging mutation burden or ploidy. **C.** Frequencies of canonical drivers in real and simulated samples. Oncogene gain-of-function, tumour suppressor loss-of-function and both types of *TP53* alterations were considered. **D.** Gene sets used for sysSVM optimisation. The training set included oncogenes (OGs) and tumour suppressor genes (TSGs), as well as *TP53*. All other damaged genes were used for prediction and assessment. These included other canonical drivers (without a proven OG or TSG role), candidate cancer genes from published cancer sequencing screens, known false positives of established driver detection methods and the remaining damaged genes. Bars indicate the number of unique damaged genes across the reference cohort of 1,000 simulated samples.

## RESULTS

### Simulation of pan-cancer datasets

In order to optimise the use of sysSVM for any cancer type, we simulated 1,000 cancer-agnostic samples starting from all TCGA tumours with matched mutation, CNV and gene expression data (Supplementary Methods, Additional File 1). We ensured that the tumour mutation and copy number burdens were similar between real and simulated samples (Figure 1B) and that gene mutation and copy number status in the simulated dataset was the same of TCGA (Supplementary Figure 1A, Additional File 1). As a result, the frequency of damaging alterations in known oncogenes and tumour suppressors was comparable between the two datasets, with *TP53*, *PIK3CA* and *CDKN2A* among the most frequently altered genes in both (Figure 1C). We further verified that gene alteration frequencies in the simulated data were not significantly biased by cancer types with large cohort sizes in TCGA (Supplementary Figure 1B, Additional File 1), confirming the suitability of the simulated data as a representative pan-cancer cohort.

The simulated cohort for sysSVM optimisation (hereafter referred to as the reference cohort) was composed of 1,000 samples with 18,455 genes damaged 309,427 times. Of these, 686 were canonical drivers with an experimentally proven role in cancer (12, 24), 1,605 were candidate cancer genes from 273 cancer screens (11), 43 were known false positive predictions of driver detection methods (1, 25) and 16,121 were the remaining damaged genes (hereafter referred to as the rest of genes; Figure 1D, Supplementary Table 2, Additional File 2). We annotated seven molecular and 25 systems-level features of all damaged genes (Supplementary Table 1, Additional File 2) and used these features for training and prediction. As a training set, we selected 457 of the 686 canonical drivers with proven roles as oncogenes (236) or tumour suppressors (221). We restricted somatic alterations of oncogenes and tumour suppressors to gain-of-function or loss-of-function alterations, respectively (Supplementary Methods, Additional File 1). Since we could not reliably define the remaining 229 damaged canonical drivers as either oncogenes or tumour suppressors, we could not restrict their somatic alterations to the appropriate type. Therefore, we did not use them for training but could still use them for prediction and performance assessment (Figure 1D), together with 43 false positives and 16,121 the rest of genes.

### sysSVM optimisation on the pan-cancer reference cohort

Using the reference cohort, we optimised sysSVM in terms of data normalisation, parameter tuning and feature selection (Figure 1A). So as not to bias the optimisation with a particular set of kernel parameters, we implemented 512 models with parameter combinations representing a sparse coverage of a standard grid search (Supplementary Note, Additional File 1). We then measured the ability of each of these 512 models to prioritise the 229 canonical drivers not used for training over the rest of damaged genes or the false positives. We did this by computing the Area Under the Curve (AUC) in each sample and taking the median AUC as representative of the whole cohort (Supplementary Methods, Additional File 1).

First, we derived the optimal settings for data normalisation in terms of centered and un-centered data (Supplementary Note, Additional File 1). All models robustly prioritised canonical drivers above the rest using either centered or un-centered data but showed lower performance in distinguishing canonical drivers from false positives (Figure 2A). We reasoned that false positives from recurrence-based driver detection methods (1) shared some features with canonical drivers. For example, they encoded long and multi-domain proteins. When removing protein length and number of domains from the feature list (Supplementary Table 2, Additional File 2), the performance substantially improved particularly for un-centred data (Figure 2B). We therefore removed protein length and number of domains from the model.

**Figure 2.**
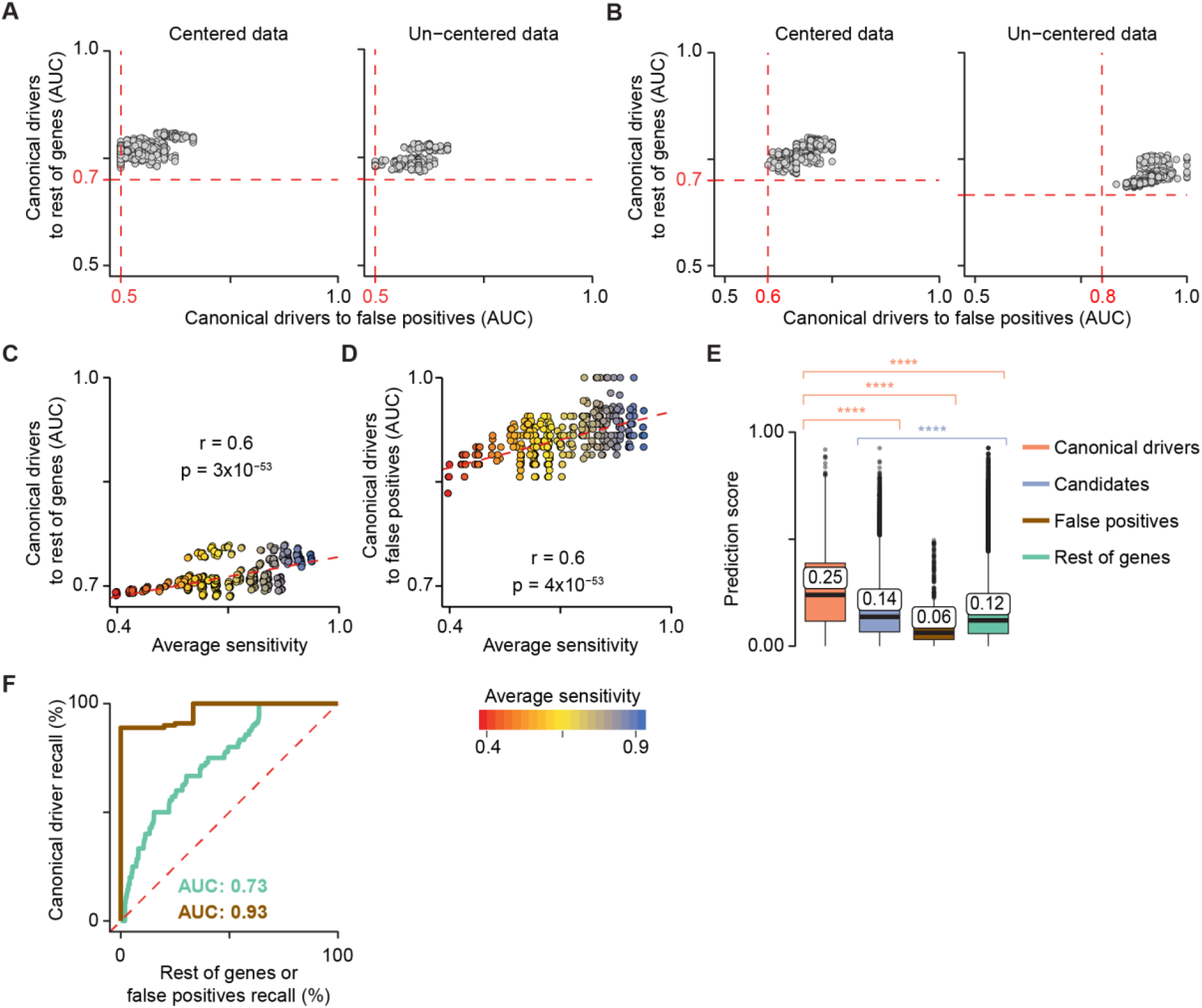
sysSVM optimisation on the simulated reference cohort. Model performances on the reference cohort using centered (left) and un-centered (right) data with all 25 systems-level features (**A**) or excluding protein length and number of protein domains (**B**). A sparse grid of 512 parameter combinations in the four kernels was tested. The performance of each model was measured using the Area Under the Curve (AUC), comparing the ranks of canonical drivers to the rest of genes and false positives. Median AUC values across all samples were plotted. Red dotted lines represent the minimum AUC values. Correlation between model average sensitivity and AUCs of canonical drivers over the rest of genes (**C**) or false positives (**D**). The sensitivity of each kernel was measured on the training set over 100 three-fold cross-validation iterations. The median values over the four kernels are plotted. R and p-values from Pearson’s correlation test are reported. Dotted red lines indicate the linear regression curves of best fit. **E.** Distributions of sysSVM2 prediction scores for different types of damaged genes in the reference cohort. Whiskers extend to 1.5 times the Inter-Quartile Range (IQR). Statistical significance was measured using two-sided Wilcoxon tests. The median values of the distributions are labelled. **** = p <2.2×10^−16^. **F.** Receiver Operating Characteristic (ROC) curves, comparing canonical drivers to the rest of genes (green) and to false positives (brown). Recall rates were calculated for each sample separately and the median ROC curve across samples was plotted. Median Areas Under the Curve (AUCs) for both comparisons are also indicated.

Second, we selected the optimal sets of parameters in each kernel. Hyper-parameter choice is known to have substantial impacts on classification and it is an open problem for one-class SVMs (26). Since the parameters for each kernel needed to be selected separately (Supplementary Note, Additional File 1), we could not use AUC of the combined multi-kernel model for assessment. Instead, we used the sensitivity of each kernel to predict canonical drivers calculated from three-fold cross-validation on the training set. Sensitivity was indeed a good predictor of the overall AUC of canonical drivers over the rest of genes (Figure 2C) and false positives (Figure 2D). We therefore developed an approach to select the parameters that conferred the highest sensitivity in multiple iterations of cross-validation (Supplementary Methods, Additional File 1). In the reference cohort, parameters chosen in this way converged within 2,000 cross-validation iterations for all kernels (Supplementary Figure 3A, Additional File 1).

Finally, since the presence of highly correlated features can hinder SVM performance (27), we performed systematic feature selection by assessing the pairwise correlations between all 25 systems-level features. Four features (gene expression in 1≤ tissues ≤6 and in ≥37 tissues; protein expression in 0≤ tissues ≤8 and central position in the protein-protein interaction network) exhibited a significant degree of inter-correlation (Pearson |r| >0.5, FDR<0.05, Supplementary Figure 3B, Additional File 1). Removing them led to faster convergence of kernel parameters (Supplementary Figure 3A, Additional File 1) and improved performance overall (Supplementary Figure 3C, Additional File 1).

Based on these results, we chose the default settings for the cancer agnostic SVM classifier, which we named sysSVM2. By default, data are un-centered but scaled to have unit standard deviation. Six of the original systems-level features are excluded resulting in a total of seven molecular and 19 systems-level features (Table 1). Finally, kernel parameters optimised on the reference cohort are provided as a default (Supplementary Figure 3A, Additional File 1), although users may perform specific cross-validation iterations on their own cohorts.

**Table 1:** Twenty-six features derived from molecular and systems-level properties of genes and used to predict cancer drivers in sysSVM2. Molecular properties describe gene alterations in individual cancer samples. Systems-level properties are global gene properties (see also Supplementary Table 1, Additional File 2). PPIN: Protein-protein interaction network. miRNA: micro RNA.

| Category | Property | Feature | Type |
| --- | --- | --- | --- |
| Molecular | Gene mutation | Mutational load (n) | Continuous |
|  |  | Non-truncating damaging mutations (n) | Continuous |
|  |  | Truncating mutations (n) | Continuous |
|  |  | Hotspot mutations (n) | Continuous |
|  | Gene copy number | Gene copy number (n) | Continuous |
|  |  | Gene is amplified | Binary |
|  |  | Gene is deleted | Binary |
| Systems-level | Gene duplication | Gene is duplicated | Binary |
|  |  | Gene is an ohnolog | Binary |
|  | Gene essentiality | Cell lines in which gene is essential (%) | Continuous |
|  |  | Gene is essential | Binary |
|  | Gene expression | Tissues expressing gene (n) | Continuous |
|  |  | Gene is expressed in 0 tissues | Binary |
| | | Gene is expressed in $7 \leq \text{tissues} \leq 36$ | Binary |
|  | Protein expression | Tissues expressing protein (n) | Continuous |
| | | Protein is expressed in $\geq 41$ tissues | Binary |
|  | Protein-protein interaction network (PPIN) | PPIN degree | Continuous |
|  |  | Protein is a PPIN hub | Binary |
|  |  | PPIN betweenness | Continuous |
|  |  | PPIN clustering coefficient | Continuous |
|  | Protein complexes | Complexes the protein is part of (n) | Continuous |
|  | miRNA interactions | miRNAs targeting the gene (n) | Continuous |
|  | Gene evolutionary origin | Pre-metazoan origin | Binary |
|  |  | Metazoan origin | Binary |
|  |  | Vertebrate origin | Binary |
|  |  | Post-vertebrate origin | Binary |

We then assessed the performance of sysSVM2 in prioritising cancer drivers over other genes. We confirmed that, overall, the prediction scores of 229 canonical drivers outside the training set were significantly higher than those of any other gene category (Figure 2E). Candidate cancer genes also scored significantly higher than the rest of genes, indicating that they were also in top ranking positions. We also measured the relative ranks of genes in individual samples using Receiver Operating Characteristic (ROC) curves. Comparing canonical drivers to the rest of genes and to false positives gave AUCs of 0.73 and 0.93, respectively (Figure 2F), demonstrating that canonical drivers were prioritised above the rest of genes and especially above false positives. This was not surprising as the properties of canonical drivers differ substantially from those of false positives (Supplementary Figure 3D, Additional File 1), further supporting that known false positives are not representative of non-cancer genes.

### Effect of training cohort size on sysSVM2 performance

The sample size of patient cohorts can highly vary across cancer types. For example, in TCGA it ranges from 32 samples for diffuse large B-cell lymphoma (DLBC) to 726 for breast cancer (BRCA, Supplementary Table 3, Additional File 2), with a median of 201 samples. We therefore sought to address how the sample size of the training cohort affected sysSVM2 performance.

Starting from all TCGA samples and using the previously described approach, we simulated 40 training cohorts, ten of which were composed of ten samples, ten of 100 samples, ten of 200 samples and ten of 1,000 samples. We then trained sysSVM2 on each of these 40 cohorts independently and used the resulting models to rank damaged genes in the reference cohort and to compare their performance.

The distributions of AUCs of canonical drivers over the rest of genes or false positives were high for all four cohort sizes (Figure 3A). This suggested that sysSVM2 was overall very effective in prioritising cancer genes independently of the training cohort size. We then compared the composition of the prioritised gene list in each sample across models of a given size. We measured a composition score of the top five genes accounting for the number and position of canonical drivers, candidate cancer genes and false positive genes (Supplementary Methods, Additional File 1). Similar to the AUC, the composition score of the top five genes was also very similar across training cohorts (Figure 3B). However, a few models trained on ten or 100 samples returned false positives in the top five positions while no false positives were predicted by models trained on larger cohorts of 200 or 1,000 samples. Finally, we measured the ratio between observed and expected canonical drivers and false positives in the top five genes (Figure 3C, Supplementary Methods, Additional File 1). Independently of the training cohort size, false positives in the top five genes were always lower than expected, confirming that sysSVM2 efficiently distinguished false positives from drivers. The number canonical drivers in the top five genes was more than twice the expected number in >85% of samples and more than five times the expected value in around 65% of samples. As with the other metrics, the performance of sysSVM2 did not change substantially with the size of the training cohort.

**Figure 3.**
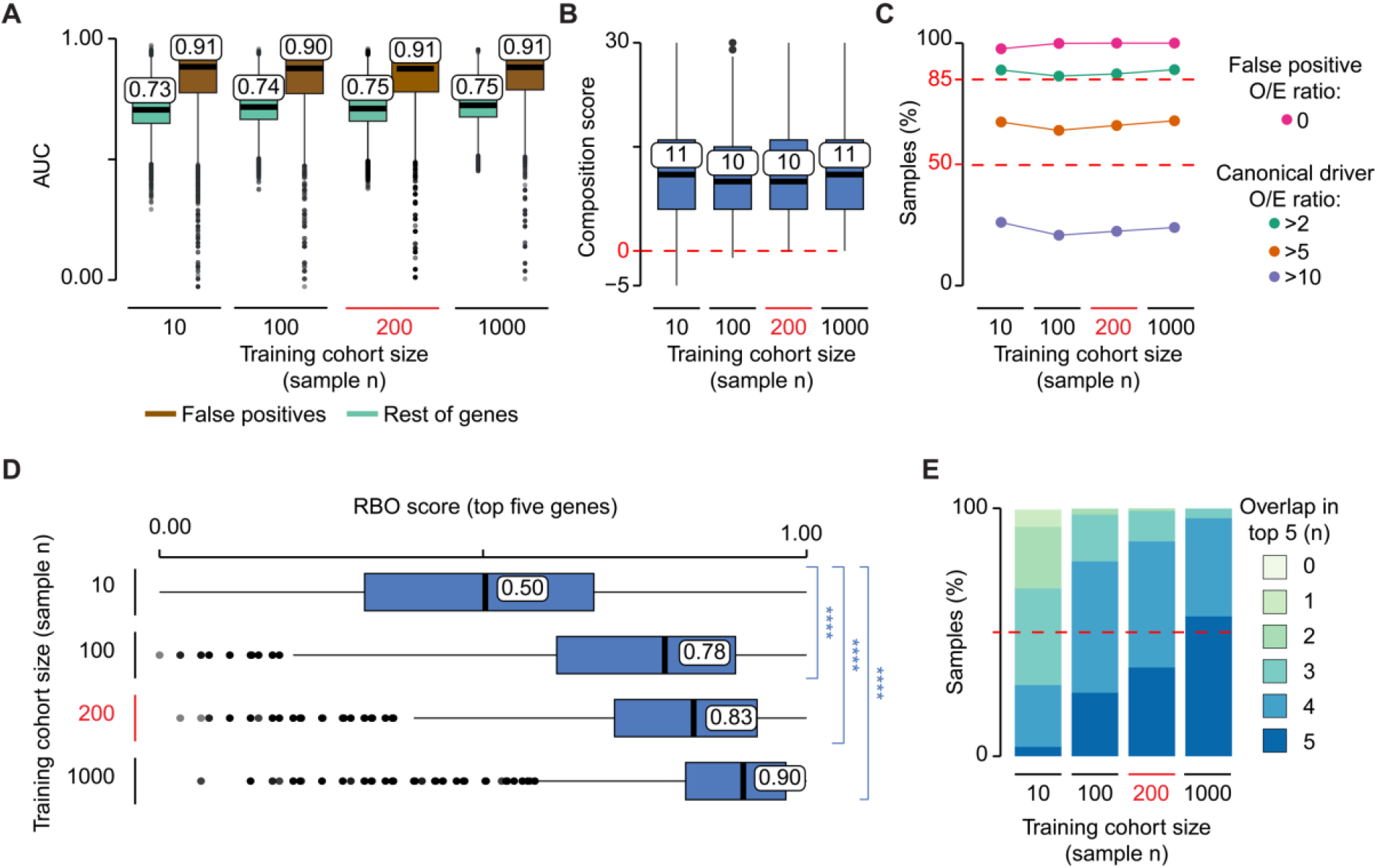
Effect of cohort size on sysSVM2 performance. **A.** Distributions of AUCs comparing the ranks of canonical drivers to the rest of genes (green) and False Positives (brown). Models were trained on ten simulated cohorts composed of ten, 100, 200 and 1,000, for a total of 40 simulated cohorts. These were then used to predict on the same reference cohort of 1,000 samples. The AUC was measured for each set of predictions in each sample. **B.** Distributions of composition scores of the top five predictions in terms of canonical drivers, candidate cancer genes, false positives and rest of genes (Supplementary Methods, Additional File 1). The composition score was measured for each set of predictions in each sample. Six training cohorts of size ten and two cohorts of size 100 gave negative composition scores in at least one sample, indicating highly ranked false positive genes. **C.** Ratios between observed and expected numbers of canonical drivers and false positives in the top five predictions (O/E ratios). For each size of the training cohort, the percentages of samples with a false positive O/E ratio of zero and canonical driver O/E ratios greater that 2, 5 and 10 are shown (Supplementary Methods, Additional File 1). **D.** Rank-Biased Overlap (RBO) score of the top five predictions in each sample (Supplementary Methods, Additional File 1). RBO scores measured the similarity between the predictions from every possible pair of models trained on cohorts of a particular sample size. Statistical significance was measured using two-sided Wilcoxon tests. **** = p <2.2×10^−16^. **E.** Distribution of the number of top five predictions shared between models trained with the same cohort size. The overlap was calculated between each pair of predictions in each sample.

Since we used the same reference cohort for prediction, we could directly compare the gene ranks in each patient across models, thus assessing their prediction stability. To do so, we measured the Rank-Biased Overlap (RBO) score that compares two ranked lists giving greater weight to the higher-ranked positions (28) (Supplementary Methods, Additional File 1). The distributions of RBO scores of the top five genes were significantly higher for large training cohorts compared to those composed of ten samples (Figure 3D). Moreover, models trained on large cohorts showed overall higher gene overlap in the top five genes (Figure 3E).

These results showed that, although sysSVM2 successfully separates canonical drivers from other genes independently of the training cohort size, small cohorts lead to occasional false positive predictions and to unstable gene ranking. Since the median cohort size of TCGA cancers is 201 samples, sysSVM2 is likely to separate canonical drivers from the rest of genes with a very low false positive rate and stable gene rankings for most cancer cohorts.

### Benchmark of sysSVM2 against existing methods

Next, we sought to compare the predictions of sysSVM2 on real cancer data to those of other driver detection methods. To do this, we used 657 Gastro-Intestinal (GI) adenocarcinomas from TCGA (73 oesophageal, 279 stomach, 219 colon and 86 rectal cancers, Supplementary Table 3, Additional File 2). Overall, this cohort had 17,122 unique damaged genes, including 438 tumour suppressors and oncogenes used for sysSVM2 training (Supplementary Table 2, Additional File 2). After ranking the remaining 16,684 damaged genes, we confirmed the overall ability of sysSVM2 to prioritise the 228 canonical drivers not used for training over the rest of damaged genes and false positives also in real data (Figure 4A).

**Figure 4.**
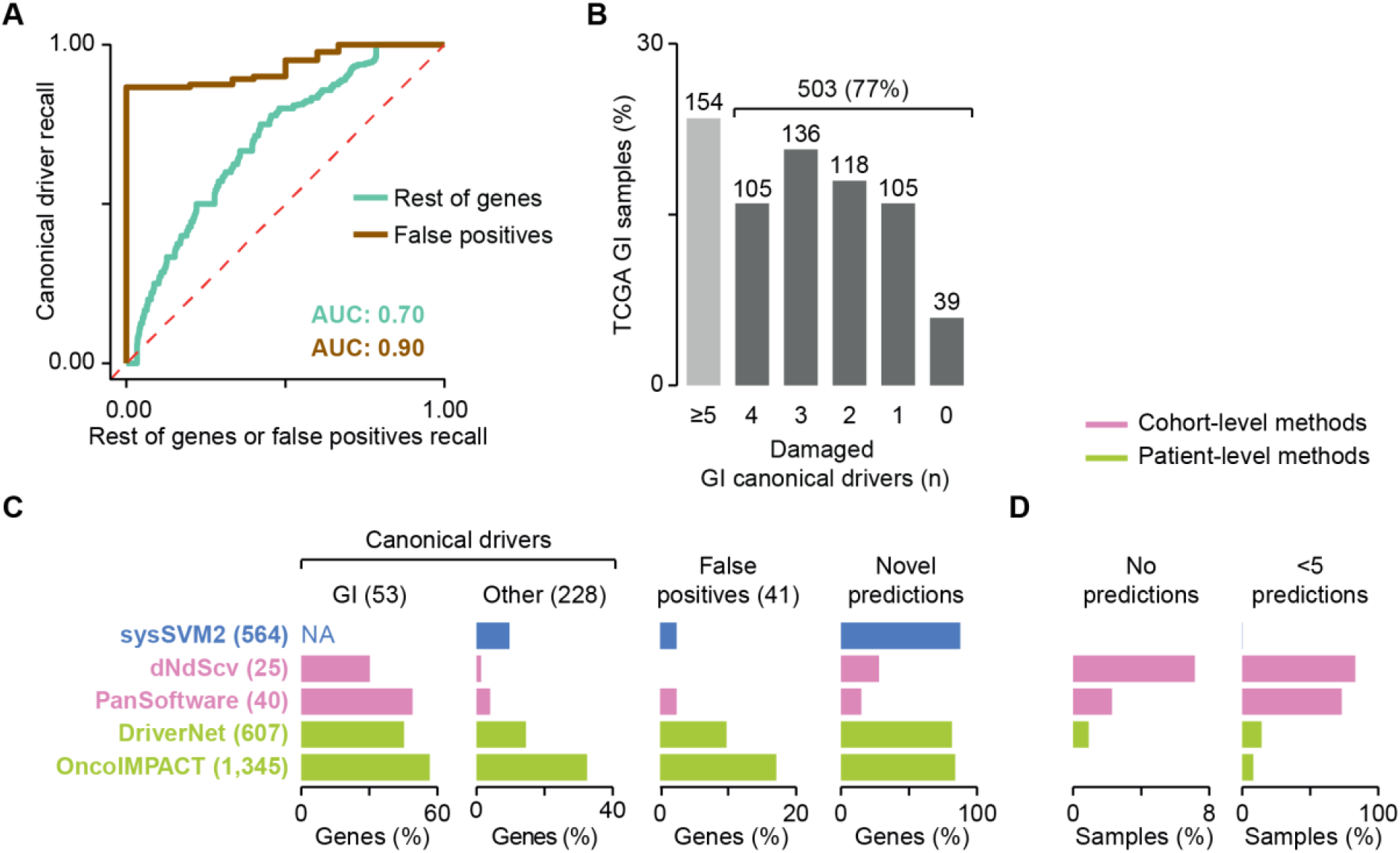
sysSVM2 benchmark on TCGA gastro-intestinal cancers. **A.** Median Receiver Operating Characteristic (ROC) curves across 657 Gastro-Intestinal (GI) samples from TCGA. Curves compare the ranks of canonical drivers to the rest of genes or to false positives. The median Areas Under the Curve (AUCs) are also indicated. **B.** Distribution of GI canonical drivers across the GI cohort. Lists of canonical drivers for each GI cancer type were obtained from NCG6 (11) and mapped to samples of the corresponding cancer type where they were damaged. Numbers of samples are indicated above each bar. Samples with five or more GI drivers did not require additional driver predictions. **C.** Comparison of performance between sysSVM2 and four other driver detection methods. The set of unique drivers predicted by each approach were compared in terms of recall of GI canonical drivers, other canonical drivers (non-GI and outside the sysSVM2 training set) and false positives and proportion of novel predictions not previously associated with a cancer driver role. The number of genes in each category is reported in brackets. The recall of GI canonical drivers could not be assessed for sysSVM2 because these were part of the training set. They were however considered as drivers by default, rather than predicted by the algorithm. NA = Not Applicable. **D.** Proportions of 657 GI samples left with no predicted drivers (left) or fewer than 5 predictions. The one sample left with fewer than 5 predictions by sysSVM2 (TCGA-FP-8210, stomach cancer) had four damaged genes overall.

To identify the list of cancer drivers of each patient, we adopted a top-up approach. Starting from the GI canonical drivers (11) damaged in each sample, we added sysSVM2 predictions progressively based on their rank to reach five drivers per patient (Supplementary Methods, Additional File 1). This was based on the assumption that each cancer requires at least five driver events to fully develop, in concordance with recent quantifications of the amount of excess mutations arising from positive selection in cancer (8, 29). While 154 patients had damaging alterations in five or more GI canonical drivers, 503 patients (77%) needed at least one prediction (Figure 4B), highlighting the need for additional cancer driver predictions. This resulted in 564 unique sysSVM2 drivers.

We then predicted the drivers in the same GI samples using two cohort-level (PanSoftware (30) and dNdScv (8)) and two patient-level (OncoIMPACT (13) and DriverNet (14)) detection methods. PanSoftware integrated 26 computational driver prediction tools and we took the list of 40 damaged drivers directly from the original publication (30), given that we used a large subset (87%) of the same TCGA GI samples. We ran the other three methods with default parameters (Supplementary Methods, Additional File 1) and obtained 25 predicted drivers with dNdScv, 607 with DriverNet and 1,345 with OncoIMPACT.

We compared sysSVM2 to the four other methods in terms of recall rates of canonical drivers or false positives, proportion of novel predictions and patient driver coverage. Overall, cohort-level methods had higher recall rates of GI canonical drivers, fewer novel predictions and a comparably low false positive recall than sysSVM2 (Figure 4C). However, unlike sysSVM2, neither cohort-level method predicted drivers in all patients, leaving the vast majority of them with less than five predictions and some with no predictions (Figure 4D).

Compared to sysSVM2, the other two patient-level methods had higher recall rates of the 228 canonical drivers, a comparably high proportion of novel predictions but higher false positive rate (Figure 4C). Namely, sysSVM2 made only one false positive prediction in one patient while DriverNet and OncoIMPACT predicted four and seven false positives in 124 and 306 patients, respectively (Supplementary Figure 4A, Additional File 1). Overall, all three methods had high patient driver coverage, but sysSVM2 outperformed the other two with only one sample where it predicted less than five drivers (Figure 4D). Interestingly, the overlap of predictions between sysSVM2 and the other patient-level methods was statistically significant (Supplementary Figure 4A, Additional File 1) even when only top-up predictions were considered (Supplementary Figure 4B, Additional File 1). This suggested that the majority of predictions converged to the same genes.

These results showed that cohort-level methods have high specificity and sensitivity to identify cancer-specific canonical drivers but often fail to find drivers in a substantial subset of patients. Compared to other patient-level detection methods, sysSVM2 outperforms them in terms of specificity and patient coverage.

### Compendium of sysSVM2 models and patient-level drivers in 34 cancer types

In order to provide a comprehensive resource of trained models and patient-level drivers, we sought to apply sysSVM2 to 7,646 TCGA samples of 34 cancer types with at least one somatically damaged gene (Supplementary Methods, Additional File 1).

To find the best training setting for the algorithm on real cancer samples, we compared the performance of sysSVM2 trained on the whole pan-cancer cohort as well as on the 34 cancer types separately. In the pan-cancer setting, we used all 477 tumour suppressors and oncogenes damaged across the whole cohort. In the cancer-specific setting, we used instead only the subsets of these genes damaged in each cancer type (Supplementary Table 3, Additional File 2). We then predicted on the remaining damaged genes and applied the top-up approach as described above, starting from the cancer-specific canonical drivers damaged in each patient (Supplementary Table 3, Additional File 2). We found that 6,067 samples (79%) required at least one sysSVM2 prediction in order to reach five drivers (Figure 5A). These corresponded to 4,369 and 4,548 unique genes in the pan-cancer and cancer-specific settings, respectively, with a significant overlap of predictions (3,896, p <2.2×10^−16^, two-sided Fisher’s exact test).

**Figure 5.**
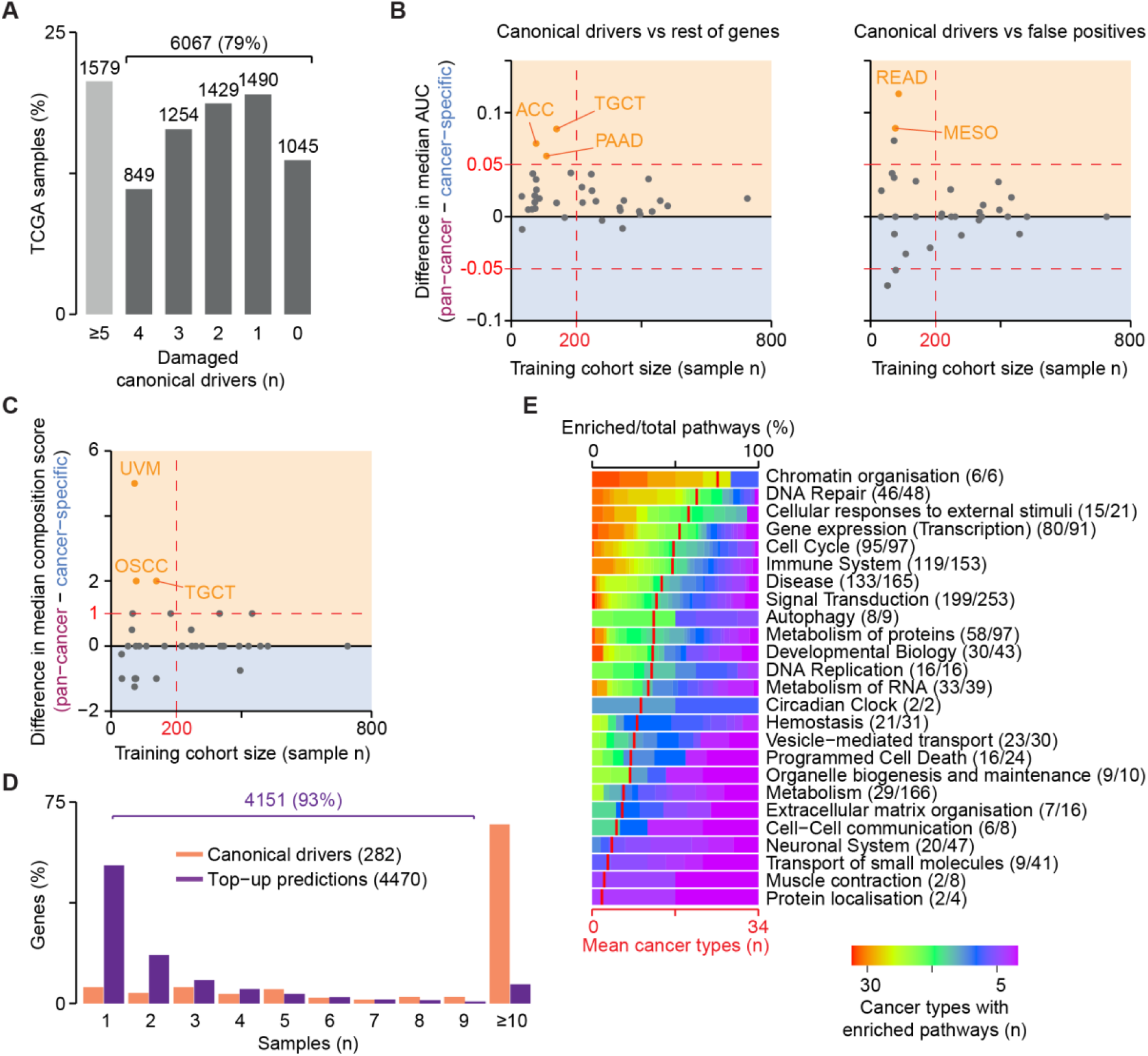
sysSVM2 predictions in 34 cancer types. **A.** Number of damaged canonical drivers per sample. Lists of canonical drivers for each cancer type were obtained from NCG (11) and mapped to samples of the corresponding cancer type. 6,067 samples with less than five canonical drivers damaged underwent the top-up procedure to reach five drivers. Difference in Areas Under the Curve (AUCs) between the pan-cancer and cancer-specific settings in ranking canonical drivers over the rest of human genes and false positives (**B**) and in the composition score of the top five predictions (**C**). The median values of the distributions in each cancer type were used for comparison, with the yellow and blue regions indicating better performance in the pan-cancer and cancer-specific settings, respectively. The number of samples used for training is indicated on the x-axis. Colour dots represent cancer types where the two settings differ both significantly (FDR <0.05, Wilcoxon rank-sum test) and substantially (|difference in medians| >0.05 for AUCs, >1 for composition score). ACC, adrenocortical carcinoma; TGCT, testicular germ cell tumours; PAAD, pancreatic adenocarcinoma; READ, rectum adenocarcinoma; MESO, mesothelioma; UVM, uveal melanoma; and OSCC, oesophageal squamous cell carcinoma. **D.** Recurrence of damaging alterations in 282 canonical driver genes and 4,470 sysSVM2 top-up predictions across 7,646 samples. **E.** Gene set enrichment analysis of sysSVM2 top-up genes, grouped in broad biological processes (Reactome level 1). Numbers of pathways enriched in at least one cancer type out of the total pathways tested are reported in brackets. Red vertical strokes indicate the mean number of cancer types that pathways from each broad process are enriched in (bottom x-axis).

We then compared the performance of pan-cancer and cancer-specific settings of sysSVM2 in prioritising canonical drivers over rest of genes or false positives. The AUCs differed significantly (FDR <0.05) and substantially (|difference in medians| >0.05) in only five cancer types (Figure 5B, Supplementary Figures 5A and 5B, Additional File 1). All of them were composed of small cohorts with <200 samples and in all cases the pan-cancer setting showed better performance than the cancer-specific setting. The composition score of the top five predictions also differed significantly and substantially (|difference in medians| >1) in only three cancer types (Figure 5C, Supplementary Figure 5C, Additional File 1). All these cancer types were again characterised by small training cohorts and showed higher performance in the pan-cancer setting. Predictions of cancer-specific models and the pan-cancer model were mostly similar, with the exception of cancer types with small training cohorts (Supplementary Figures 5D and 5E, Additional File 1). Overall, these results confirmed the trend observed in the simulated data and indicated that the pan-cancer and cancer-specific settings performed similarly well in most cases, except for small cohorts where the pan-cancer model performed better.

Based on these results, we used the pan-cancer setting for cancer types with small cohorts (N <200) and the cancer-specific setting for the others, as this could reflect cancer-type specific biology without jeopardising performance or stability. The final list of patient-specific predictions in 34 cancer types was composed of 4,470 unique genes, the vast majority of which (93%) were rare (<10 patients) or patient-specific (Figure 5D, Supplementary Table 4, Additional File 2). A gene set enrichment analysis on these genes revealed 984 enriched pathways overall (Reactome level 2 or above, FDR <0.01, Supplementary Methods, Additional File 1, Supplementary Table 5, Additional File 2). Interestingly, when mapping these pathways to broader biological processes (Reactome level 1), a few processes were widely enriched in almost all cancer types (Figure 5E). These included well-known cancer-related processes such as chromatin organisation (31), DNA repair (32), cell cycle (33) and signal transduction (34). Therefore, although not recurring across patients, sysSVM2 predictions converged to perturb similar biological processes that are known to contribute to cancer.

### sysSVM2 predictions in an independent cancer cohort

We finally sought to assess whether the sysSVM2 models trained on TCGA could be applied for driver prediction in a cancer type not included in TCGA. We therefore analysed 36 osteosarcomas from the Pan-Cancer Analysis of Whole Genomes (PCAWG) consortium (29). Osteosarcoma is a rare, genetically heterogeneous bone cancer with poor prognosis and only six well-established canonical drivers (35, 36).

We annotated the genomic data of the PCAWG cohort finding 4,969 damaged genes overall with a median of 93 damaged genes per sample (Supplementary Table 2, Additional File 2). Only two of these samples had three damaged osteosarcoma canonical drivers while 19 (53%) of them had no canonical driver (Figure 6A), highlighting the need for further predictions. Given the small cohort size, we used the TCGA pan-cancer setting to rank the damaged genes in each osteosarcoma. Considering the top five predictions per sample, we got 129 unique genes (Supplementary Table 6, Additional File 2), which were poorly recurrent across samples (Figure 6B), reflecting again the genetic heterogeneity of osteosarcoma.

**Figure 6.**
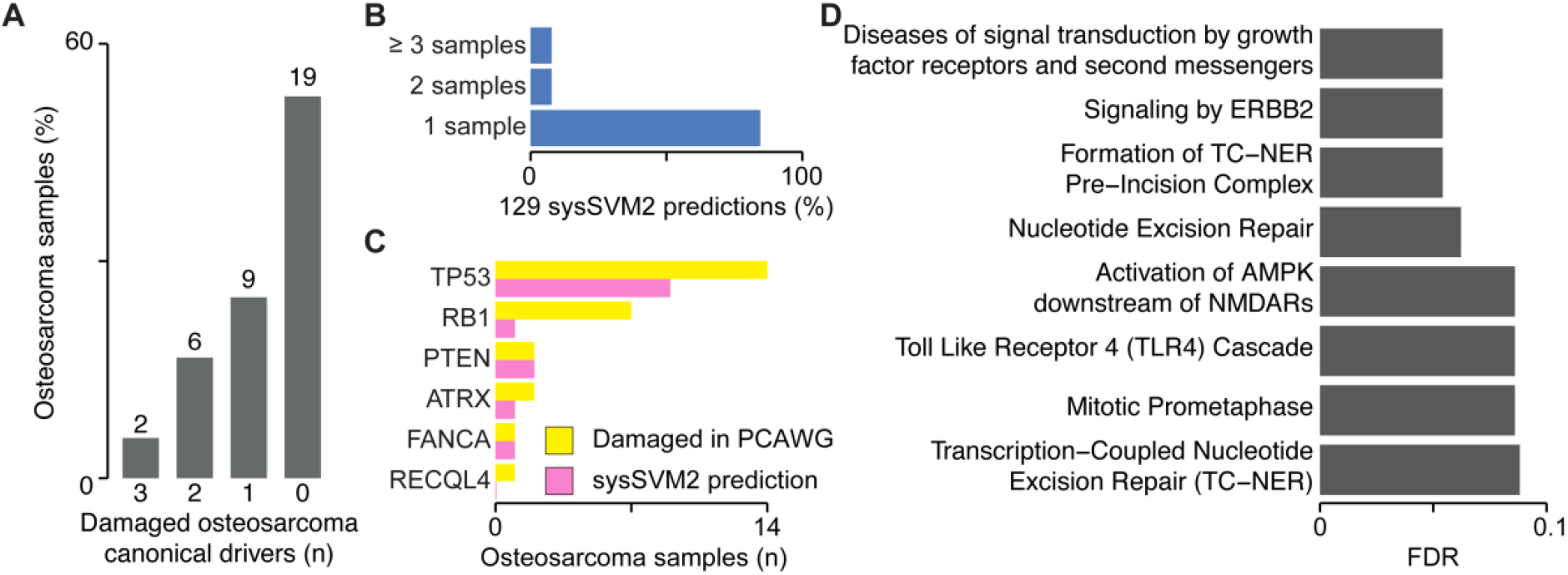
Validation of sysSVM2 in osteosarcoma. **A.** Distribution of osteosarcoma canonical drivers across the PCAWG osteosarcoma cohort. Lists of canonical drivers for osteosarcoma derived from the literature (35, 36) and mapped to samples where they were damaged. Numbers of samples are indicated above each bar. **B.** Recurrence of the 129 sysSVM2 predictions across the PCAWG osteosarcoma cohort. The percentages of genes that are predicted in 1, 2 and ≥3 are shown. **C.** Patient-level predictions of osteosarcoma canonical drivers by sysSVM2 when considering the top five genes. The number of samples in which each canonical driver is damaged (yellow) and predicted as a driver by sysSVM2 (pink) is shown. **D.** Gene set enrichment analysis of 81 sysSVM2 predictions with no previously reported involvement in cancer. Reactome level 2 and above were considered and pathways with FDR <0.1 are shown.

At the cohort level, sysSVM2 predictions included five of the six (83%) osteosarcoma canonical drivers (35, 36). At the patient level, the six osteosarcoma canonical drivers were damaged 27 times and in 14 of these cases (53%) they were in the top five predictions (Figure 6C). This proportion rose to 81% when considering the top ten predictions. In addition to osteosarcoma canonical drivers, 26 sysSVM2 predictions were canonical drivers in other cancer types, 16 were candidate cancer driver genes and 81 had no previously known involvement in cancer (Supplementary Table 6). Despite this, these 81 genes were enriched in eight pathways (FDR <0.1), most of which have a known role in cancer (Figure 6D). Moreover, they included genes known to promote osteogenesis such as *YAP1* and *YES1* (37, 38).

These results showed that sysSVM2 is able to identify reliable cancer drivers in individual patients even for cancer types not used for training. This has relevant implications particularly in the case of rare cancers that are poorly studied and have little genomic data available.

## DISCUSSION

Identifying the complete repertoire of driver events in each cancer patient holds great potential for furthering the molecular understanding of cancer and ultimately for precision oncology. While many recurrent driver genes have now been identified, the highly heterogeneous long tail of rare drivers still poses great challenges for detection, validation and therapeutic intervention.

Our method allows to identify driver genes in individual patients. These genes converged to well-known cancer-related biological processes and further studies could potentially use these predictions to investigate particular aspects of cancer biology, such as driver clonality and their progressive acquisition during cancer evolution. Extending the algorithm with additional sources of data is another avenue for future work. For example, transcriptomic and epigenomic data could enhance the ability of sysSVM2 to identify driver events. Additionally, recent efforts have identified a large number of driver events in non-coding genomic elements (29). Given such a training set of true positives, sysSVM2 could be further developed to identify non-coding drivers in individual patients, as long as appropriate features could be identified. The general approach of identifying drivers using a combination of molecular and systems-level properties affords great flexibility for such developments.

It is increasingly common for sequencing studies to integrate multiple tools for driver detection (30), since building a consensus can make results robust to the weaknesses of individual methods. sysSVM2 also has its weaknesses. For example, while systems-level properties distinguish cancer genes as a set, there are some cancer genes that do not follow this trend (11) and are thus likely to be missed by the algorithm. Our approach in the current work of topping up known driver genes with predictions from sysSVM2 is a simple example of how sysSVM2 can be used in conjunction with other approaches. More broadly, it is likely the case that patient-level driver detection will eventually rely on an entire ecosystem of different methods. In this work, we have demonstrated that there is a place for sysSVM2 in such an ecosystem.

## CONCLUSIONS

In this work, we developed a cancer-agnostic algorithm, sysSVM2, for identifying cancer driver in cancer individual patients. By refining the machine learning approach upon which the original algorithm was built (18), we broadened its applicability to the pan-cancer range of malignancies represented in TCGA. sysSVM2 successfully and stably prioritises canonical driver genes for most publicly available cancer cohorts. For those composed of fewer samples, the models optimised on the whole pan-cancer dataset offer a valid alternative. Moreover, compared to other patient-level driver detection methods, sysSVM2 has better patient coverage and a particularly low rate of predicting established false positives. sysSVM2 can be used to identify driver alterations in individual patients and rare cancer types where canonical drivers are insufficient to explain the onset of disease, as we have validated in osteosarcoma. This potentially opens up further research and therapeutic opportunities.

## AVAILABILITY AND REQUIREMENTS

**Project name:** sysSVM2

**Project home page:** https://github.com/ciccalab/sysSVM2

**Operating system:** Platform independent

**Programming language:** R

**Other requirements:** R version greater than 3.5

**License:** CRICK Non-commercial License Agreement v2.0

**Any restrictions to use by non-academics:** Commercial use will require a license from the rights-holder. For further information contact.

## LIST OF ABBREVIATIONS

SNV: Single nucleotide polymorphism
Indel: Insertion or deletion
CNV: Copy number variant
SVM: Support vector machine
TCGA: The Cancer Genome Atlas
AUC: Area under the curve
ROC: Receiver operating characteristic
DLBC: Diffuse large B-cell lymphoma
BRCA: Breast cancer
RBO: Rank-biased overlap
GI: Gastro-intestinal
FDR: False discovery rate
PPIN: Protein-protein interaction network

## DECLARATIONS

### Availability of data and materials

Platform-independent R code to implement sysSVM2, along with a README file and an example dataset, is available at https://github.com/ciccalab/sysSVM2. The recommended settings as described in this manuscript are set as default values. However, users can modify many aspects of the implementation, including selection of features, data normalisation and kernel parameters. Models trained in pan-cancer and cancer-specific settings in 34 TCGA cancer types are also provided. This software code is protected by copyright. No permission is required from the rights-holder for non-commercial research uses. Commercial use will require a license from the rights-holder. For further information contact.

Original data for annotating systems-level properties were obtained from the following publicly available sources:

BioGRID: https://thebiogrid.org/
CORUM: http://mips.helmholtz-muenchen.de/corum/
DIP: http://dip.doe-mbi.ucla.edu/dip/Main.cgi
EggNOG: http://eggnogdb.embl.de/#/app/home
GTEx: https://www.gtexportal.org/home/
HPRD: http://www.hprd.org/
MIntAct: https://www.ebi.ac.uk/intact/
miRecrods: http://c1.accurascience.com/miRecords/
miRTarBase: http://mirtarbase.mbc.nctu.edu.tw/php/index.php
OGEE: http://ogee.medgenius.info/browse/
PICKLES: https://hartlab.shinyapps.io/pickles/
Protein Atlas: https://www.proteinatlas.org/
Reactome: https://reactome.org/
RefSeq: https://www.ncbi.nlm.nih.gov/refseq/

## Authors’ contributions

F.D.C. conceived and directed the study. J.N. performed data simulation and algorithm assessment and optimisation under the supervision of F.D.C. and C.Y. H.M. carried out the pan-cancer TCGA analysis with the help of J.N. F.D.C. and J.N. wrote the manuscript. All authors approved the manuscript.

## Acknowledgements

The results published here are in whole or part based upon data generated by The Cancer Genome Atlas managed by the NCI and NHGRI. We thank Damjan Temelkovski for testing sysSVM2.

## ADDITIONAL FILES

**Additional File 1:** Supplementary Note, Supplementary Methods, Supplementary Figures (DOCX). sysSVM2 rationale and algorithm description. Algorithm implementation and assessment. Figure S1 Comparison of simulated and TCGA samples. Figure S2 Selection of binary features derived for PPIN and tissue expression properties. Figure S3 Parameter convergence and feature selection. Figure S4 Patient-level comparison of driver detection methods. Figure S5 Setting comparison for sysSVM2 training on TCGA data.

**Additional File 2:** Supplementary tables (XLSX). Table S1 Features of genes used in sysSVM2. Table S2 Cohorts and genes used in the study. Table S3 Application of sysSVM2 to TCGA samples. Table S4 Driver predictions in 7,646 TCGA samples. Table S5 Gene set enrichment analysis of TCGA predictions. Table S6 sysSVM2 driver predictions in PCAWG osteosarcomas.

