## Additional File 1 for "Pan-cancer detection of driver genes at the single-patient resolution"

**Supplementary Note.** sysSVM rationale and algorithm description
**Supplementary Methods.** Algorithm implementation and assessment

**Supplementary Figure 1.** Comparison of simulated and TCGA samples
**Supplementary Figure 2.** Selection of binary features for PPIN and tissue expression properties
**Supplementary Figure 3.** Parameter convergence and feature selection
**Supplementary Figure 4.** Patient-level comparison of driver detection methods
**Supplementary Figure 5.** Setting comparison for sysSVM2 training on TCGA data

**Supplementary Table 1**. Features of genes used in sysSVM2
**Supplementary Table 2.** Cohorts and genes used in the study
**Supplementary Table 3.** Application of sysSVM2 to TCGA samples
**Supplementary Table 4.** Driver predictions in 7,646 TCGA samples
**Supplementary Table 5.** Gene set enrichment analysis of TCGA predictions
**Supplementary Table 6.** sysSVM2 driver predictions in PCAWG osteosarcomas

**Supplementary Note.** sysSVM rationale and algorithm description

### sysSVM rationale

The central principle of sysSVM is that cancer genes differ from other human genes by an array of properties that can be used to prioritise new cancer genes in individual samples (1). These properties fall into two categories, molecular properties and systems-level properties. Both categories are encoded as continuous or binary features of a Support Vector Machine (SVM) classifier.

Molecular properties describe the somatic alterations that genes undergo in cancer tumours (sequence and copy number alterations). Molecular properties change in each analysed cohort because they will derive from the patient cancer sequencing data. The molecular properties used in optimising sysSVM for pan-cancer use are encoded by five continuous features (total exonic mutational load, non-truncating damaging mutations, truncating mutations, hotspot mutations, gene copy number) and two binary features that indicate whether the gene is amplified or deleted. Details on how damaging mutations, gene amplifications and gene deletions are derived from cancer sequencing data are provided in Supplementary Methods, Additional File 1.

Systems-level properties are global properties of genes that are not directly related to cancer but nevertheless differentiate cancer genes from the rest of human genes. Some systems-level properties specifically describe tumour suppressor genes and oncogenes, allowing sysSVM to match the gain- or loss-of-function alterations with genes that are more similar to oncogenes and tumour suppressor genes, respectively. For example, canonical drivers, and in particular tumour suppressor genes, are less likely than other human genes to have duplicated loci elsewhere in the genome (2, 3. On the other hand, oncogenes are more likely to have evolved through whole-genome duplication events that occurred at the basis of vertebrates (i.e. to be ohnologs) {D'Antonio, 2011 #25). Canonical drivers are essential gene more often and in a higher proportion of cell lines than other genes (4). They are expressed in a wider range of healthy human tissues, both at the gene and protein levels. Proteins encoded by canonical drivers have distinct topological features in the protein-protein interaction network (PPIN), namely they have higher PPIN degree, betweenness and clustering coefficients than other proteins (3). Canonical driver proteins also participate in more complexes (5) and their genes are targeted by more miRNAs (2), suggesting that the expression of cancer genes is tightly regulated. Tumour suppressor genes tend to be old genes with a pre-metazoan origin, while oncogenes tend to originate in metazoans and cancer drivers in general are depleted in post-vertebrate genes (2). Finally, cancer genes encode longer proteins with more domains than other human genes (5, 6).

Systems-level properties are encoded in sysSVM as either continuous or binary features, depending on their nature. To account for the fact that some SVM kernels learn more efficiently from binary features (1) and to more comprehensively describe the underlying properties, additional binary features were derived for PPIN properties as well as for gene and protein expression. For PPIN features, hubs and central proteins were defined as those in the top 25% of degree and betweenness distributions, respectively. These thresholds were chosen because they marked out the high tails of the continuous distributions while describing sufficiently large numbers of proteins to be informative (Supplementary Figures 2A,B). For gene and protein expression, the distributions of tissues expressing the gene/protein are broken down into four distinct sections (Supplementary Figures C,D). For protein expression, two binary features (9-34 tissues, and 35-40 tissues) were not statistically different between cancer proteins and the rest of proteins. All systems-level features considered for the optimisation of sysSVM for pan-cancer use differed significantly either between canonical drivers, oncogenes or tumour suppressor genes and the rest of genes. This resulted in 25 systems-level features (Supplementary Table 1, Additional File 2), which underwent further feature selection to obtain the 19 final features used in sysSVM2 (Table 1).

**The sysSVM algorithm**

As previously described (1), sysSVM consists of four one-class Support Vector Machines (7) (SVMs) trained on the molecular and systems-level properties of the canonical cancer drivers damaged in a cohort of patients. It then ranks damaged genes outside the training set based on how similar their properties are to those of canonical cancer drivers.

The four one-class SVMs use different kernels, which control how each one learns from the training set. A kernel $k$ measures how similar the features of two genes are. Let $x$ and $y$ denote the features of two genes. Then an SVM measures their similarity as $k(x, y)$. The kernels used in sysSVM are:

- Linear: $k(x, y) = x \cdot y$
- Polynomial: $k(x, y) = (x \cdot y)^d$
- Radial: $k(x, y) = \exp -\gamma|x - y|^2$
- Sigmoid: $k(x, y) = \tanh\,(\gamma\, x \cdot y)$

Here $x \cdot y$ denotes the standard dot product, $d$ and $\gamma$ are parameters, and $\tanh$ is the hyperbolic tangent function. sysSVM combines the outputs of the four kernels into a single score to use for prediction. The sysSVM algorithm consists of three stages: feature mapping; model selection; and training and prediction.

**Step 1. Feature mapping**

In feature mapping, molecular and systems-level properties are mapped to the damaged genes of the training cohort. These include seven molecular features (relating to mutation and copy number status) and 19 systems-level features. An additional six systems-level features are available but are excluded from the model by default, since their inclusion was seen to worsen performance. The full list of features is given in Supplementary Table 1, Additional File 2.

**Step 2. Model selection**

Model parameters are then selected. The four one-class SVMs are controlled by certain parameters, and a grid search is implemented to select the best parameters for each kernel separately. These parameters and their default grid ranges are:

- Nu ($\nu$, all kernels): represents an upper bound on the proportion of the training set that can be classed as outliers. Values range from 0.05 to 0.35 in steps of 0.05.
- Gamma ($\gamma$, radial and sigmoid kernels): controls the level of influence of individual training points on the model. Values assessed are $\gamma = 2^x$, where $x \in \{-7, -6, \ldots, 4\}$
- Degree ($d$, polynomial kernel): the degree of the polynomial kernel function, chosen from the set $\{3, 4, 5\}$.

In the first implementation of sysSVM (1) an additional parameter was tuned: $\gamma$ in the polynomial kernel. In this setting, the polynomial kernel had the form $k(x, y) = (\gamma\, x \cdot y)^d$. However, in this case $\gamma$ simply controls an overall scaling of the kernel, since $(\gamma\, x \cdot y)^d = \gamma^d (x \cdot y)^d$. Constant scalings such as this do not change the behaviour of

SVMs, and so the $\gamma$ parameter is redundant. Thus, in sysSVM2 $\gamma$ is fixed to 1 for the polynomial kernel by default.

The selection of the best parameter values for each kernel is described in the Supplementary Methods, Additional File 1. Cross-validation iterations are performed on the training set, and at each iteration the sensitivity of each parameter combination is assessed. The final parameters are chosen on the basis of having a high average and low standard deviation of sensitivity.

Tuning the parameters for each kernel separately, rather than together, greatly reduces the number of combinations to be assessed. For example, using the default ranges above, the linear kernel has 7 parameter combinations, the radial and sigmoid kernels have $7 \times 12 = 84$ combinations each, and the polynomial kernel has $7 \times 3 = 21$ combinations. This gives a total of $7 + 84 + 84 + 21 = 196$ parameter combinations to assess. By contrast, using the same parameter grid and tuning parameters based on the combined performance of the full sysSVM model would give $7 \times 84 \times 84 \times 21 = 1{,}037{,}232$ parameter combinations to assess. Thus, the parameters for each kernel must be selected separately for the grid search to be tractable.

**Step 3. Training and prediction**

Once the parameters for each kernel have been selected, the four SVMs are trained using the entire training set of canonical drivers.

The trained sysSVM model can then be used for prediction in individual samples. To combine the outputs of the four kernels, a combined score $S_{gs}$ is calculated for each gene $g$ in sample $s$. $S_{gs}$ measures the similarity of the features of gene $g$ to those of the training set. It combines the rank of $g$ in sample $s$ according to each of the four kernels, in such a way that the final score is normalised between 0 and 1. High ranks in each kernel are given exponential weighting and the kernels are weighted according to their sensitivity, with more sensitive kernels contributing more to the score. If $R_{kgs}$ is the rank of $g$ in sample $s$ according to the decision value of kernel $k$, $N_s$ is the total number of damaged genes in sample $s$ and $\mu_k$ is the mean sensitivity of kernel $k$ as assessed by cross-validation iterations, then the score is

$$S_{gs} = \frac{\sum_{k=1}^{4}\left(-\log_{10}\left(\frac{R_{kgs}}{N_s}\right) \times \mu_k\right)}{4 \times \log_{10}(N_s)}$$

#### Parameter ranges for the grid search

When measuring the performance of various implementations of sysSVM using the Area Under the Curve (AUC) as in Figure 2, we used a reduced grid search range. This assessment was based on the performance of the full model combining all four kernels, and thus the full range would have been prohibitively costly ($1{,}037{,}232$ parameter combinations, as described above). Instead, we considered $\nu \in \{0.05, 2\}$, $\gamma = 2^x$ where $x \in \{-7, 3, 0, 4\}$, and $d \in \{2, 4\}$. This resulted in a total of $(2) \times (2 \times 4) \times (2 \times 4) \times (2 \times 2) = 512$ parameter combinations to assess, where the brackets enclose the number of parameters for the linear, radial, sigmoid and polynomial kernels, respectively. We chose these ranges to provide a sparse coverage of the parameter grid used in the standard model selection step of sysSVM.

#### Data normalisation and the one-class Support Vector Machine

In this section we discuss the theory behind why data centering is not optimal for the one-class Support Vector Machines in sysSVM.

The one-class SVM algorithm of Schölkopf *et al.*(7) aims to construct a decision boundary which encloses the region of input space $\mathcal{X}$ where the training set lies. It then classes points inside the decision boundary as similar to the training set (cancer genes), and points outside as outliers (non-cancer genes). In order to draw complicated decision boundaries efficiently, it uses a kernel function $k$ to (implicitly) map the training set to a feature space, $\mathcal{F}$. In $\mathcal{F}$, the algorithm aims to separate the training set from the origin, using a linear decision boundary (a hyperplane). The final decision boundary is the result of mapping this hyperplane back to the input space $\mathcal{X}$.

The problem with data centering is most clear for the linear kernel. With this choice of kernel, feature space $\mathcal{F}$ corresponds exactly to input space $\mathcal{X}$. If data are centered around the origin (*i.e.* zero), then the algorithm attempts to separate the data from their own centre, which is clearly inappropriate. Indeed, experiments with a toy example indicate that this can lead to extreme sensitivity to small changes in the training set (data not shown).

On the other hand, centering is not an issue for the radial kernel, since it is translationally invariant: if gene features $x$ and $y$ are shifted by some constant value $c$, then it can be seen from the radial kernel function that $k(x-c, y-c) = k(x, y)$.

For the polynomial and sigmoid kernels, the effect of centering is less clear. They both preserve the origin – that is, the origin in $\mathcal{X}$ corresponds to the origin in $\mathcal{F}$. This can be seen from the fact that $k(0, x) = 0$ for any set of gene features $x$, with either kernel. This might suggest that centering should be avoided with these kernels, and this is further borne out in toy model experiments.

**Supplementary Methods.** Algorithm implementation and assessment

### Pan-cancer TCGA data annotation and simulation

Sequence mutations (SNVs and indels) for 9,079 samples were obtained from the MC3 release of TCGA (8) and annotated with ANNOVAR (9) (downloaded April 2018) and dbNSFP v3.0 (10). Only mutations identified as exonic or splicing were retained. Damaging mutations included (1) truncating (stopgain, stoploss, frameshift) mutations; (2) missense mutations predicted by at least five out of seven functional prediction methods (SIFT (11), PolyPhen-2 HDIV (12), PolyPhen-2 HVAR, MutationTaster (13), MutationAssessor (14), LRT (15) and FATHMM (16)) , at least two out of three conservation-based methods (PhyloP (17), GERP++RS (18) and SiPhy (19)); (3) splicing mutations predicted by one of two splicing-specific methods (ADA (10) and RF) and (4) hotspot mutations identified by OncodriveCLUST (20) v1.0.0.

Copy number data (array intensities) for 11,379 samples were obtained from the Genomic Data Commons portal (https://portal.gdc.cancer.gov/). Copy Number Variant (CNV) segments, sample ploidy and sample purity values were obtained using ASCAT (21) v2.5.2, with 9,873 samples passing quality control. Segments were intersected with the exonic coordinates of 19,549 human genes that were derived as previously described (4) in the reference genome hg38, and converted to hg19 coordinates using the UCSC liftOver tool (http://genome.ucsc.edu/). Genes were considered to have undergone a CNV if at least 25% of their transcribed length was covered by a segment. RNA-Seq data were used to filter out false positive CNVs. Fragments per kilobase million (FPKM) values were obtained for 10,974 samples and only samples with matched copy number and expression data were retained. Damaging CNVs included homozygous gene losses (copy number 0 and FPKM <1 over mean cancer type purity) and gene amplifications (copy number >2 x sample ploidy).

Considering only one sample per patient, a total of 7,630 samples had matched mutation, CNV and RNA-Seq data that passed ASCAT quality controls and had at least one damaged gene. These comprised 535,615 damaging mutations and 2,041,598 damaging CNVs in 18,784 genes (Supplementary Table 2, Additional File 2).

To simulate samples that reproduced the molecular features of real TCGA samples, the damaging mutation burden and ploidy were measured for the whole TCGA cohort. Then, 1,000 random combinations of damaging mutation burden and ploidy were extracted and assigned to the simulated samples. To preserve the same frequency of damaging alterations, damaged genes were extracted from within TCGA samples that had similar values of damaging mutation burden (+/-10%, for mutations) and ploidy (+/-0.1, for CNVs) and assigned to the simulated samples. Overall, the final simulated dataset of 1,000 samples contained 69,269 damaging mutations and 252,409 damaging CNVs in 18,455 genes.

As a training set, 220 tumour suppressors with 2,433 loss-of-function alterations (truncating mutations, missense or splicing damaging mutations, homozygous deletions, or double hits) and 236 oncogenes with 5,539 gain-of-function alterations (hotspot mutations, missense or splicing damaging mutations or gene amplifications) were retained. For *TP53* both loss- and gain-of-function alterations (n=352) were considered to be driver events and included in the training set. 301,103 damaging alterations in the remaining 17,998 human genes not present in the training set were used for prediction (Supplementary Table 2, Additional File 2). Somatic alterations in tumour suppressors and oncogenes that were not of the appropriate types (i.e. gain-of-function in tumour suppressors and loss-of-function in oncogenes) were discarded from further analysis.

#### Annotation of systems-level properties

Systems-level properties of human genes were obtained from various sources. Duplicated gene loci (genes with 60% protein sequence shared with another locus) were identified as previously described (4), and ohnolog status was obtained from Nakatani *et al.* (22). Data on gene essentiality in cell lines were obtained from the PICKLES (September 2017) (23) and OGEE v2 (24) databases. Genes in PICKLES were considered to be essential in a cell line if they had Bayes factor >3, while original annotations of essentiality from OGEE were retained. mRNA expression data for healthy human tissues were downloaded from GTEx v7 (25) and the Protein Atlas v18 (26), and a gene was considered to be expressed in a tissue if median TPM >1 in both databases (or in one database if that tissue was not included in both). Categorical protein expression data (indicating whether or not a protein was expressed) for healthy human tissues was taken from the Protein Atlas v18 (26). The protein-protein interaction network was constructed as previously described (4), from the union of BioGRID v3.4.157 (27), MIntAct v4.2.10 (28), DIP (February 2018) (29) and HPRD v9 (30), resulting in 16,322 proteins and 289,368 interactions supported by at least one original publication. Topological properties (PPIN degree, betweenness and centrality) were calculated using custom scripts. Genes were identified as participating in complexes using data taken from CORUM (July 2017) (31), HPRD v9 (30) and Reactome v63 (32). The number of miRNAs targeting a gene was calculated using data from miRTarBase v7.0 (33) and miRecords v4.0 (34). The evolutionary origin (age) of genes was identified as previously described (2), using data from EggNOG v4.5.1 (35). Protein length was obtained from RefSeq (36) release 94, and the number of protein domains were obtained from InterPro (37) annotations in UniProt (38) release 2019_07. From these properties, 25 systems-level features were derived to be used for classification, of which 19 were retained after feature selection (Supplementary Table 1, Additional File 2).

For each feature, missing values were imputed using the median or mode (for numeric and categorical features, respectively) of available data for canonical drivers and the rest of genes separately. All features used in the model significantly differentiated cancer drivers from other human genes (Supplementary Table 1, Additional File 2).

**Kernel parameter tuning**

sysSVM2 uses four one-class Support Vector Machines (SVMs), with linear, radial, sigmoid and polynomial kernels following three steps: feature mapping, model selection, and training and prediction (Figure 1A, Supplementary Note, Additional File 1). sysSVM2 is implemented in R (39), and SVM training and prediction are carried out using the e1071 (40) package.

Model selection was updated from the original sysSVM formulation to improve convergence. A parameter grid search was carried out for each kernel separately, for a total of 196 kernel-parameter combinations (Supplementary Note, Additional File 1). The aim was to select parameters that resulted in a model with a high sensitivity and stability (low standard deviation of sensitivity). The model sensitivity for each parameter combination was assessed on the simulated training set using three-fold cross-validation for 5,000 iterations. For each kernel $k$ and parameter combination $i$, the mean $\mu_{ki}$ and standard deviation $\sigma_{ki}$ of the sensitivity were calculated across the cross-validation iterations. These were then converted into z-scores $z_{ki}^{(\mu)}$ and $z_{ki}^{(\sigma)}$, which measured the relative values of mean and standard deviation between the different parameter combinations such that:

$$\sum_i z_{ki}^{(\mu)} = \sum_i z_{ki}^{(\sigma)} = 0$$

and

$$\text{Variance}_i\left(z_{ki}^{(\mu)}\right) = \text{Variance}_i\left(z_{ki}^{(\sigma)}\right) = 1.$$

Finally, we defined the $\Delta_z$ score as:

$$\Delta_z = z_{ki}^{(\mu)} - z_{ki}^{(\sigma)}$$

High $\Delta_z$ scores corresponded to parameter combinations that had high mean sensitivity and low standard deviation relative to the other combinations for that kernel. The four parameter combinations (one per kernel) with the highest $\Delta_z$ scores were selected and used to train the four kernels on the entire training set.

**Performance assessment**

The performances of all driver prediction models tested in sysSVM2 were measured using five metrics: Area Under the Curve (AUC); composition score; Observed/Expected (O/E) ratios; Rank-Biased Overlap (RBO) score and overlap of the top five predictions between models.

For the AUC, Receiver Operating Characteristic (ROC) curves were derived for each sample individually by comparing the ranks of canonical drivers not used for training to known false positive genes and to the rest of human genes. For both of these comparisons, the median AUC was then measured across samples. The median ROC curve across the cohort was also derived by calculating the median true positive rate for each value of a false positive rate.

The composition score assessed the top five predictions in each sample, and measured the prevalence and ranks of different types of genes. The score *S* was calculated as a weighted sum according to the following formula:

$$S = \sum_{g=1}^{5} w_g \times t_g.$$

The weight $w_g$ of each gene *g* in the top five was such that higher-ranked genes were assigned greater weight. Specifically, $w_g = 6 - r_g$ where $r_g$ is the rank of gene $g$ (with 1 being the highest). The type contribution $t_g$ of gene g was defined for different gene categories as follows: cancer-specific canonical drivers ($t_g = 3$); other canonical drivers (2); cancer-specific candidate genes (1.5); other candidate cancer genes (1); false positives ($-1$); other genes (0).

Ratios between observed and expected numbers of canonical drivers and false positives in the top five predictions (O/E ratios) were calculated as follows:

$$\text{Canonical driver O/E ratio} = \frac{\text{Canonical drivers in top 5 genes}}{\text{Total canonical drivers in sample}} \times \frac{\text{Total damaged genes in sample}}{5}$$

$$\text{False positive O/E ratio} = \frac{\text{False positives in top 5 genes}}{\text{Total false positives in sample}} \times \frac{\text{Total damaged genes in sample}}{5}$$

These formulas accounted for the fact that the difficulty of ranking canonical drivers (or false positives) in the top five predictions depends on how many damaged genes there are in each sample. The percentages of samples where the observed number of canonical drivers in the top five predictions was higher than twice, five and ten times the expected numbers, and where the observed number of false positives was lower than expected, were calculated.

The RBO score (41) was used to assess the similarity of the top five predictions from pairs of models. It measures the overlap of ranked lists at incrementally increasing depths using a convergent series. Including a correction for finite lists of length five, it was calculated according to the formula:

$$\text{RBO} = \frac{1-p}{1-p^5}\sum_{d=1}^{5} p^{d-1} \times A_d.$$

where *p* was set to 0.9 (41), *d* indicates depth in the rankings (starting from the top-ranked elements), and $A_d$ is the overlap of the two lists, restricted to depth *d*.

**TCGA sample analysis**

Starting from the TCGA samples with matched mutation, copy number and RNA-Seq data, more stringent filters were applied for homozygous deletions and gene amplifications. Genes with CN=0 that either (1) had one or more mutations in their sequence, or (2) were expressed at greater than 1 FPKM over sample purity (as opposed to mean cancer type purity) were re-annotated as having CN=1. Amplifications were filtered by applying Wilcoxon tests to compare expression levels between amplified and non-amplified cases, for each gene separately. Only genes for which amplification was associated with overexpression at FDR <0.05 were retained as having damaging amplifications. The resulting dataset comprised 7,651 samples at least one damaging alteration. To prevent sysSVM2 training sets from being biased by individual samples, five samples each accounting for more than 50% of the damaged genes in their respective cancer types were removed, leaving 7,646 samples for further analysis. For the purposes of measuring performance and overlap of the cancer-specific and pan-cancer sysSVM2 settings, 752 samples with fewer than ten damaged genes were removed.

To tune the kernel parameters of sysSVM2, 10,000 three-fold cross validations were run in each cancer type. In cases where parameters did not converge, additional 15,000 cross-validations were run only for the non-converging parameters, with all other parameters fixed at their converged values. This was done for gamma of the sigmoid kernel in adrenocortical carcinoma (ACC), oesophageal adenocarcinoma (OAC), colon adenocarcinoma (COAD), kidney renal clear cell carcinoma (KIRC) and stomach adenocarcinoma (STAD), as well as for the degree parameter of the polynomial kernel in lung squamous cell carcinoma (LUSC). Prediction were assessed using the same metrics as for simulated data (AUC, composition score, RBO score and overlap of top five predictions).

Lists of cancer driver genes in individual TCGA samples were identified using a top-up procedure as follows. First, a list of canonical driver genes for each of the 34 cancer types in TCGA was obtained from NCG (4). Cancer-specific canonical drivers damaged in each sample were considered as cancer drivers for that sample. In samples with five or more such drivers, no further prediction was done. Otherwise, the highest-ranked genes from sysSVM2 were added so that there were five drivers in total. Samples with five or fewer damaged genes overall were not considered for the pan-cancer analysis. For the purpose of comparing sysSVM2 to other driver detection methods on gastro-intestinal cancer data one sample with less than five damaged genes (TCGA-FP-8210, stomach cancer) was included for completeness.

sysSVM2 predictions in 657 gastro-intestinal (GI) cancer samples were compared to those of PanSoftware (8); dNdScv (42); OncoIMPACT (43); and DriverNet (44). Forty genes identified as GI cancer drivers by PanSoftware was taken from the original publication (8). These genes were considered as drivers in every sample in which they were damaged. dNdScv was implemented with default parameters, taking all mutations as input. Genes were considered to exhibit significant signs of positive selection if they had a FDR <0.05. OncoIMPACT was implemented with default parameters, using the gene network provided. Nonsilent mutations were used as input for mutations. Copy number amplifications were filtered as described above, and copy number losses included heterozygous and homozygous deletions. In each sample, all genes that shared the highest score were taken as the predicted drivers. DriverNet was implemented with default parameters, using the same gene network and data inputs as for OncoIMPACT. The events identified in each sample were taken as the predicted drivers.

Human pathways for gene set enrichment analysis were obtained from Reactome v72 (45). Before testing, pathways were restricted to those of level 2 or higher, and with between 10 and 500 genes (total of 1,429 pathways containing 10,178 genes). In each cancer type separately, the unique set of top-up predictions from sysSVM2 was tested for enrichment in pathways containing at least one prediction, using one-sided hypergeometric tests The resulting p-values across all cancer types were corrected for False Discovery Rate (FDR) using the Benjamini-Hochberg method (46), as a single set.

### Annotation of PCAWG osteosarcoma data

Variant call vcf files (SNVs, indels) and bed files (copy number segments) were downloaded for 36 PCAWG osteosarcoma samples from the ICGC Data Portal (https://dcc.icgc.org/). Variants were annotated as described for TCGA samples, except for gene amplifications that could not be filtered based on overexpression since matched gene expression data were unavailable. Instead, a more stringent threshold of copy number >2.5 x sample ploidy was used. This resulted in a total of 4,969 damaged genes across the cohort, comprising 3,270 unique genes (Supplementary Table 2, Additional File 2).

**Supplementary Figure 1.** Comparison of simulated and TCGA samples

**A**

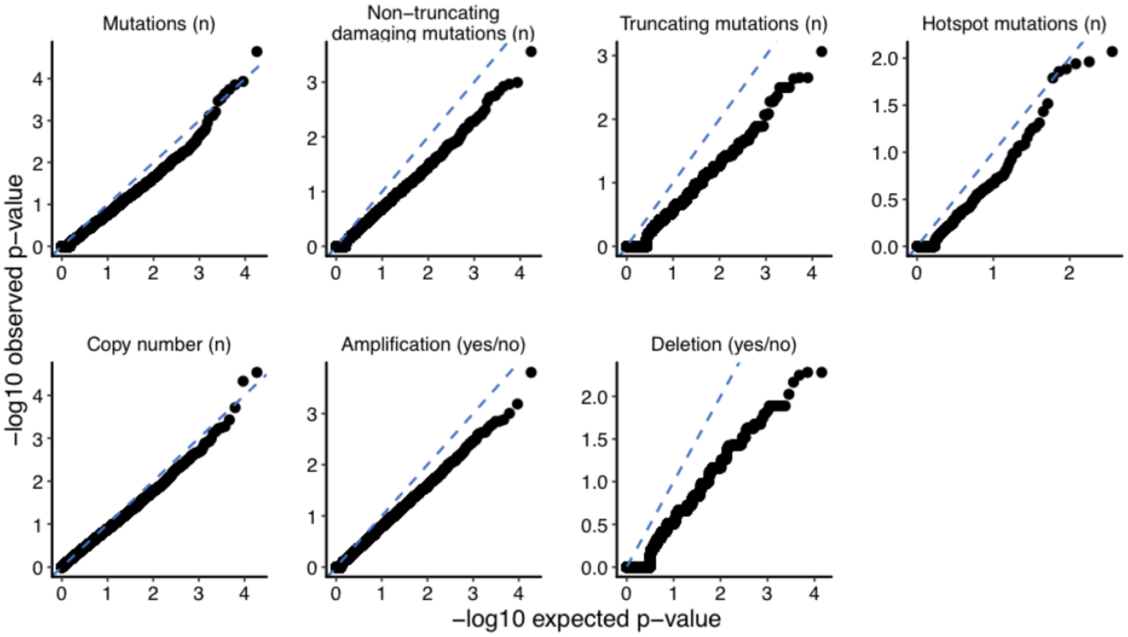


**B**

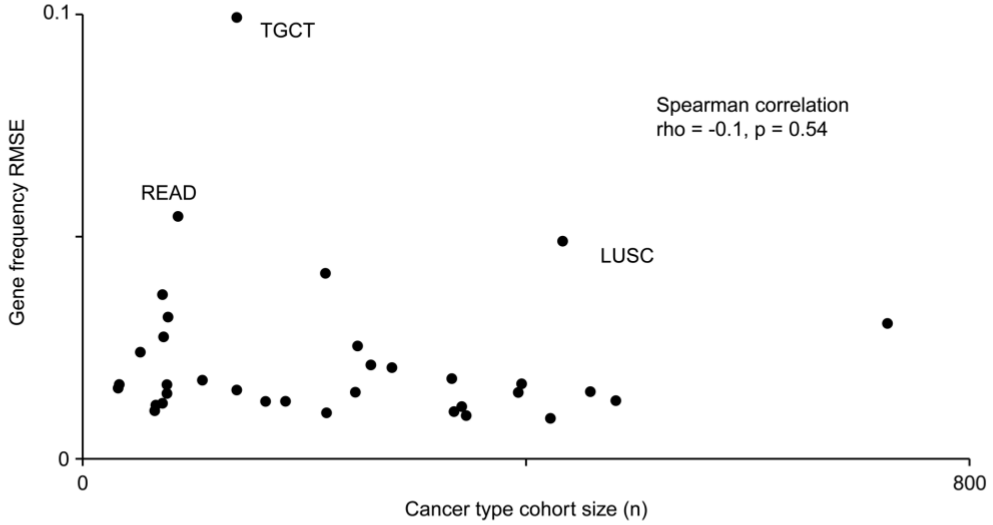


**A.** Quantile-quantile (QQ) plots illustrating comparisons of molecular features of genes between TCGA (n=7,630) and simulated reference (n=1,000) cohorts. Features were compared using Poisson tests (mutation features), Wilcoxon tests (copy number) and Fisher tests (amplifications and deletions).

**B.** For each cancer type in TCGA, the gene alteration frequency profile was calculated as the proportion of samples with a damaging alteration in each gene. This was then compared to the gene alteration frequency profile of the simulated reference cohort of 1,000 samples using the root mean-squared error (RMSE). This was calculated as $RMSE = \sqrt{\frac{1}{19549}\sum_{g=1}^{19549}\left(f_g^{TCGA} - f_g^{sim}\right)^2}$, where $f_g^{TCGA}$ and $f_g^{sim}$ are the frequencies of gene g being damaged in the TCGA and simulated cohorts, and 19549 is the total number of human genes(4). Higher values of the RMSE indicate greater differences between the simulated reference cohort and the TCGA cohort for that cancer type. Cancer types with RMSE >0.05 are labelled. There was no significant correlation between cohort size and RMSE (Spearman correlation p=0.54, rho=-0.11).

**Supplementary Figure 2.** Selection of binary features for PPIN and tissue expression properties

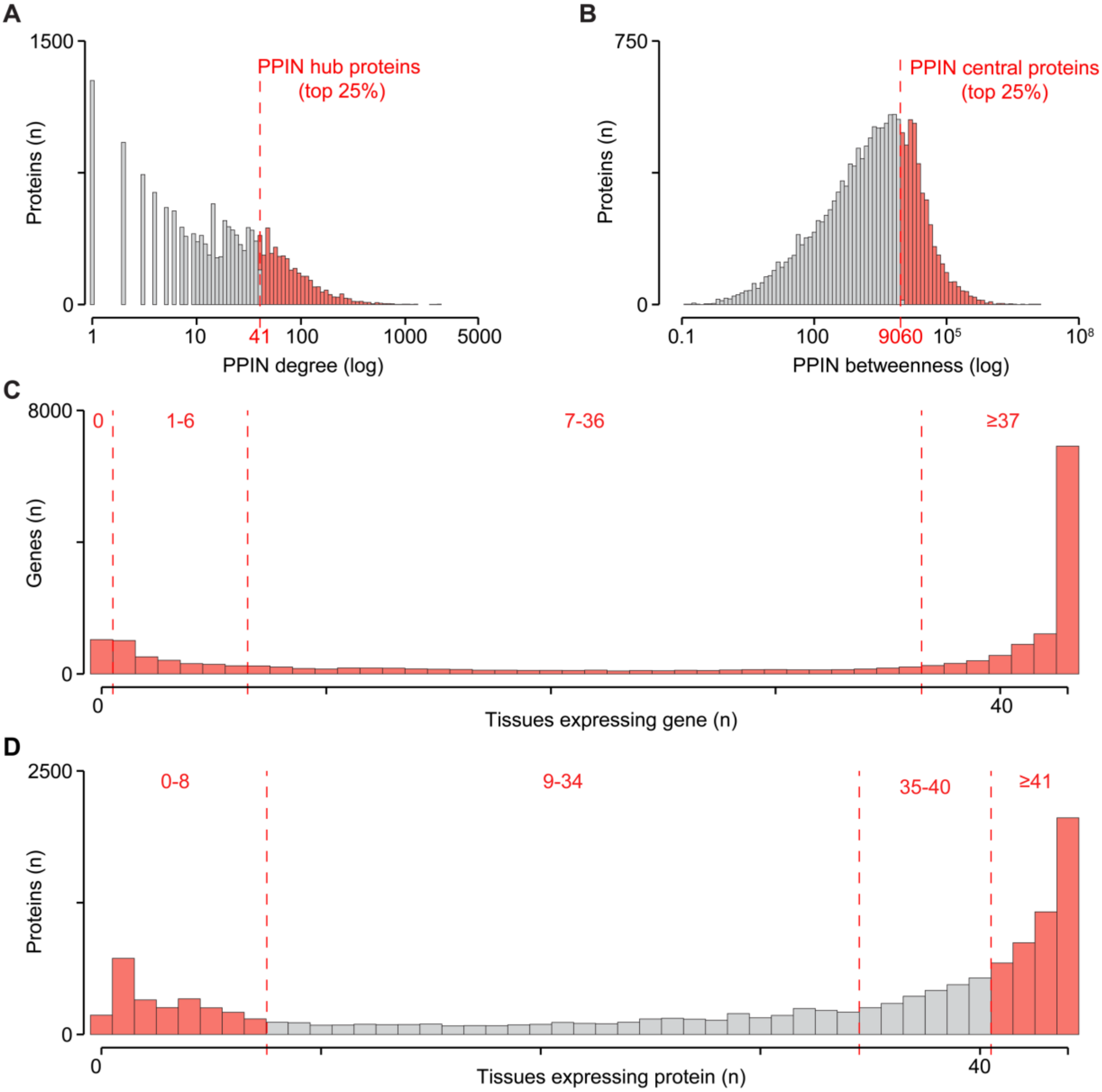


Distributions of protein-protein interaction network (PPIN) degree (**A**) and betweenness (**B**) for 16,322 human proteins. Proteins in the top 25% for degree and betweenness values were designated as hubs and central proteins, respectively.

Distributions of the number of tissues expressing 18,641 genes (**C**) and 13,001 proteins (**D**). Both distributions were divided into four sections based on their shape to derive binary expression features.

Features highlighted in red were statistically different between canonical drivers and the rest of genes (Supplementary Table 1, Additional File 2).

**Supplementary Figure 3.** Parameter convergence and feature selection

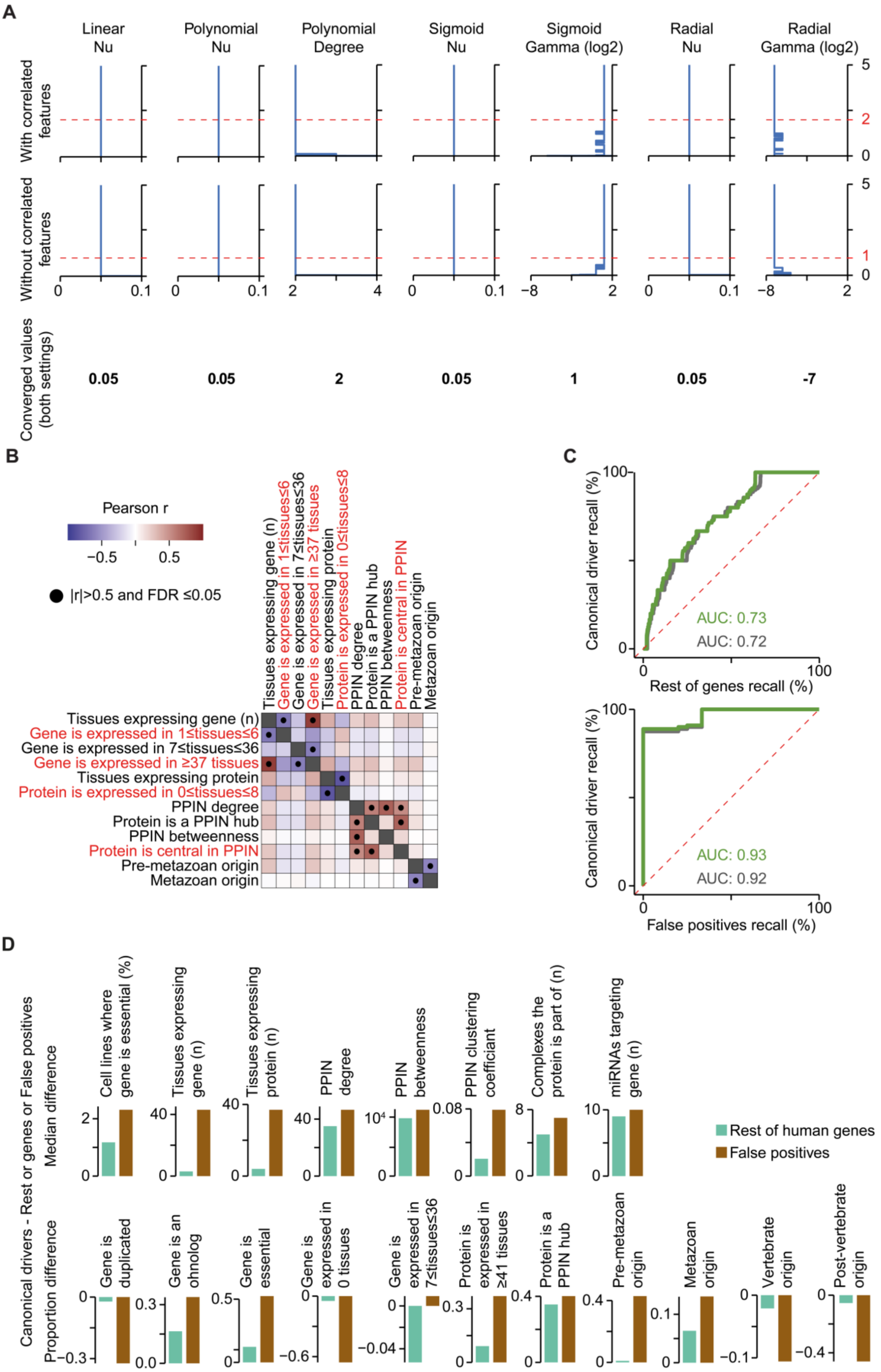

**A.** Parameter convergence on simulated data, with (top) and without (middle) correlated features. The four kernels had a total of seven parameters chosen from a grid search. Cumulative parameter selections are indicated over a series of 5,000 cross-validation iterations, but parameter choices converged within 2,000 iterations. The final selected parameter values are indicated at the bottom and were the same for both settings (with and without correlated features).

**B.** Correlation between systems-level features. Pearson correlation coefficient was measured between all possible pairs of features reported in Supplementary Table 1, Additional File 2 considering all genes with available data. Only features with at least one significant correlation (Pearson $|r|>0.5$, FDR<0.05) are shown. Three correlated feature pairs (Protein-Protein Interaction Network (PPIN) degree – PPIN hub, PPIN degree – PPIN betweenness, and pre-metazoan and metazoan origin) were retained because they were complementary in their description of gene properties. In particular, PPIN betweenness is a distinct topological property from degree, and hubs included proteins with a large range of degrees (41 to 2,116). The evolutionary origin features instead described the two most common epochs for the origin of human genes (11,624 pre-metazoan, 3,076 metazoan, Supplementary Table 1, Additional File 2). Final features removed are in red.

**C.** Comparison of model performances with (green) and without (grey) correlated features. Median Receiver Operating Characteristic (ROC) curves across samples are plotted, comparing the ranks of canonical drivers to the rest of genes (top) and to false positives (bottom). The median Areas Under the Curve (AUCs) are also indicated. The green ROC curves are the same as those shown in Figure 2F.

**D.** Difference in average features between canonical drivers and the rest of human genes (green) and false positives (brown). For each of the three gene sets and each of the 19 systems-level features selected for sysSVM2, either the median value (continuous features, top row) or the proportion of genes for which the feature was positive (binary features, bottom row) was calculated. The difference in these values between canonical drivers and both the rest of genes and false positives is shown for each feature. For all features except Gene is expressed in 7≤tissues≤36, the difference between canonical drivers and false positives is greater than the difference between canonical drivers and rest of genes, while the direction of the difference is the same.

**Supplementary Figure 4.** Patient-level comparison of driver detection methods

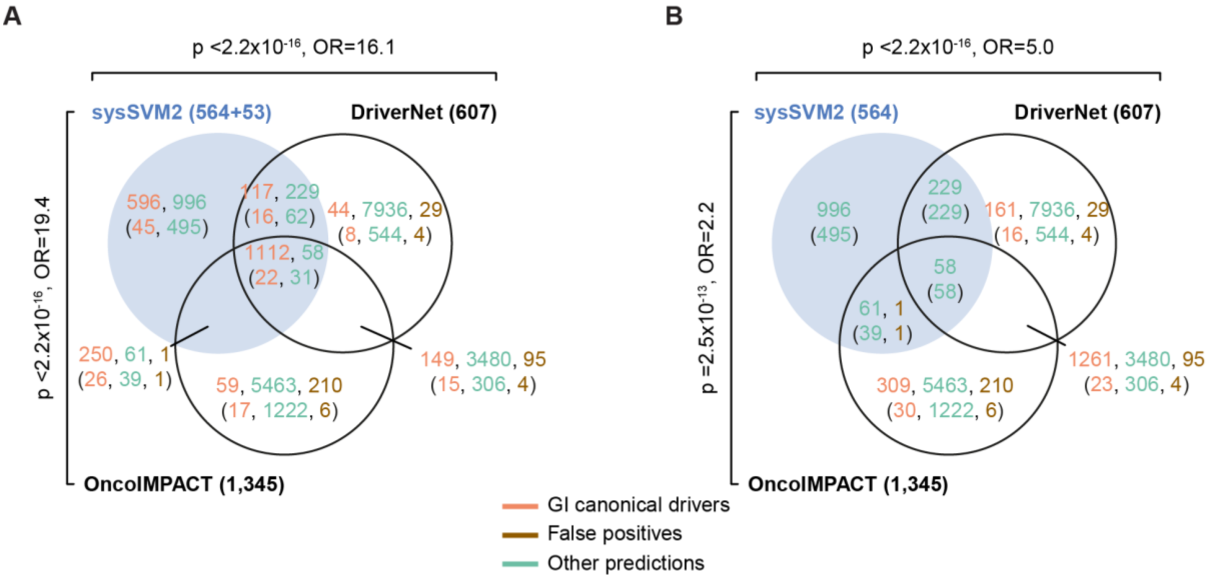


Overlap of driver predictions in individual samples, between sysSVM2, DriverNet[3] and OncoIMPACT[4]. The sysSVM2 predictions were considered both with (**A**) and without (**B**) the GI canonical drivers. The numbers of overlapping patient-level predictions are indicated, along with the number of unique genes in each set in brackets. P-values and Odds Ratios (ORs) for pairwise overlap were calculated using Fisher's exact test, taking into account all damaged genes in all samples.

**Supplementary Figure 5.** Setting comparison for sysSVM2 training on TCGA data

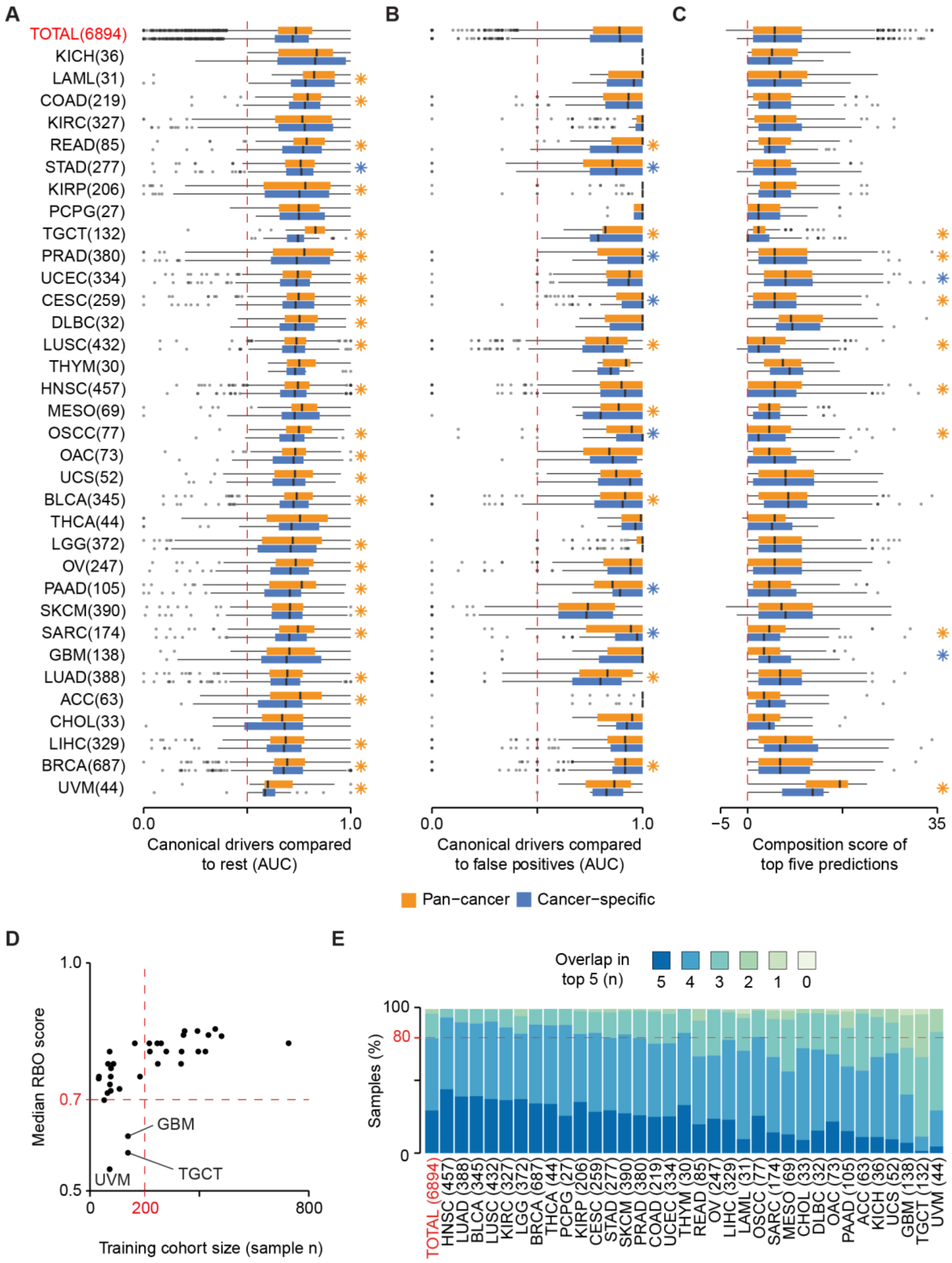


Distributions of performance of the pan-cancer and cancer-specific settings across samples measured through (**A**) AUC of the rank of canonical drivers over the rest of human genes (**B**) AUC of the rank of canonical drivers over false positive genes and false positive genes and

(**C**) the composition score of the top five predicted genes (Supplementary Methods, Additional File 1). Median values across samples are indicated. * FDR <0.05. Yellow stars indicate better performance of the pan-cancer model, blue stars indicate better performance of the cancer-specific model.

**D.** Rank-Biased Overlap (RBO) scores measuring the similarity between the top five predictions in the pan-cancer and cancer-specific settings. Median RBO scores for each cancer type are shown on the y-axis, and the number of samples used for training is shown on the x-axis. UVM, uveal melanoma; GBM, glioblastoma multiforme; TGCT, testicular germ cell tumours.

**E.** Overlap in the top five predictions between the pan-cancer and cancer-specific settings, for each cancer type. The numbers of samples whose predictions were considered are indicated in brackets.
